# α-synuclein pathology drives multi-scale reorganization of fronto-limbic networks and cognitive flexibility in the common marmoset

**DOI:** 10.64898/2026.06.11.731687

**Authors:** Alessandro Zanini, Jesleen Saini, Jiayue Yang, Audrey Dureux, Maëva Gacoin, Constance Dollet, Kevin D. Johnston, Wen Luo, Irina Shlaifer, Thomas M. Durcan, Philippe Huot, Sriram Jayabal, Timothy J. Bussey, Lisa M. Saksida, Vania Prado, Marco A.M. Prado, Ravi S. Menon, Justine C. Cléry, Stefan Everling

## Abstract

Understanding how α-synuclein misfolding and spreading affect brain systems is central to synucleinopathy research, yet evidence from primate models remains limited. Here, we combined histology, awake structural MRI, awake resting-state fMRI, regional homogeneity, actimetry and touchscreen behavioral testing to longitudinally track the consequences of striatal α-synuclein seeding in the common marmoset. Phosphorylated α-synuclein inclusions were detected from 2 months post-injection and spread progressively to distributed cortical and subcortical regions bilaterally, accompanied by regionally specific structural atrophy. Functional imaging revealed widespread disruption of large-scale networks, particularly within fronto-limbic circuits, alongside reductions in local functional coherence. Despite the absence of overt motor or sleep deficits, animals exhibited a selective impairment in cognitive flexibility, with preserved learning and task engagement. These findings indicate that striatal α-synuclein seeding induces a multi-scale reorganization of brain networks preferentially affecting circuits supporting cognitive flexibility and limbic processing, establishing a primate platform for studying early stages of synucleinopathy.

## INTRODUCTION

Synucleinopathies, including Parkinson’s disease (PD), dementia with Lewy bodies (DLB), and multiple system atrophy, are defined by the progressive misfolding and intracellular accumulation of α-synuclein (αSyn) into oligomeric and fibrillar assemblies that disrupt synaptic, mitochondrial, and proteostatic function^1,2^. A defining feature of these disorders is the spatial propagation of protein misfolding and aggregation beyond its site of origin: misfolded and abnormally phosphorylated αSyn (at S129 residue) spreads across anatomically connected regions, correlating with the sequential emergence of motor, cognitive, autonomic, and sleep disturbances across disease stages^3–5^.

Converging evidence supports a prion-like mechanism of propagation, whereby misfolded αSyn serves as a template to drive further aggregation in connected, vulnerable neuronal populations^6^. This framework repositions synucleinopathies as progressive network disorders rather than regionally circumscribed lesion syndromes^7^. Yet despite extensive histological characterization of propagation, how molecular pathology translates into large-scale alterations in brain structure, functional connectivity, and behaviour over time remains poorly understood particularly in primates^8,9^, a gap that limits both mechanistic insight and the development of circuit-level therapeutic targets for humans.

Rodent preformed fibril (PFF) models have established core principles of αSyn spread, demonstrating transmission from striatal injection sites to interconnected cortical and limbic regions^10,11^. However, the translational relevance of these models is constrained by fundamental differences in brain organization and connectivity relative to humans. Non-human primates offer a substantially closer approximation to human cortico-basal ganglia-thalamo-cortical and limbic architecture^12,13^. Among them, the common marmoset (*Callithrix jacchus*) is particularly well suited for longitudinal systems-level investigation: it possesses a differentiated neocortex, shares high sequence identity with human αSyn, and affords tractable experimental access for repeated *in vivo* preclinical high-field magnetic resonance imaging (MRI) and behavioural assessment^14–16^.

Prior work has established that intracerebral injection of α-synuclein fibrils into the marmoset striatum induces progressive phosphorylated α-synuclein pathology, dopaminergic neurodegeneration, and Lewy body–like inclusions, confirming the feasibility of primate PFF models and their histopathological relevance to synucleinopathy^15^. However, these studies have remained anchored to molecular and neurochemical endpoints, leaving open a fundamental question: how does α-synuclein propagation reshape large-scale brain organization over time, and what are its functional and behavioural consequences? No study to date has tracked the longitudinal evolution of α-synuclein pathology alongside structural integrity, functional network dynamics, and cognition within a single non-human primate cohort. In particular, whether striatal seeding produces early, selective cognitive deficits prior to and independent of overt motor symptoms has not been tested in a primate model, despite compelling evidence for this sequence in rodents^17^. Addressing this gap is critical: the relationship between pathological spread, circuit-level dysfunction, and the emergence of specific behavioural phenotypes holds the key to understanding why different synucleinopathies produce distinct clinical profiles, and to identifying early, network-level biomarkers of disease progression.

Here, we injected marmoset-derived αSyn PFF into the caudate and putamen and tracked the consequences of αSyn seeding longitudinally using histology, structural MRI, resting-state functional MRI (fMRI), regional homogeneity analyses, actimetry, and cognitive behavioural testing (see Supplementary Figure 1 for detailed timeline). We show that fibril seeding initiates a progressive, multi-scale cascade from localized inclusion pathology to distributed structural atrophy, large-scale network disruption, and selective impairment of cognitive flexibility in the absence of overt motor deficits. These findings establish a systems-level account of circuit vulnerability in synucleinopathy and provide a non-human primate platform to improve translation to humans in early stages of the disease.

## RESULTS

### Immunohistochemistry

Immunofluorescence revealed widespread distribution of pS129-positive αSyn pathology following unilateral PFF injection in the caudate and putamen. At the earliest timepoint examined (2 months post-injection), pS129-immunoreactivity was detectable but spatially restricted to the injection sites (caudate and putamen), substantia nigra, hippocampal region and, in a lesser extent, frontal, orbitofrontal, and anterior cingulate areas including 9, 11, 14 and 32. By subsequent timepoints (5, 7, 11, and 13 months), pathology had expanded progressively both ipsilaterally and contralaterally, with pS129-positive inclusions observed across a broad rostrocaudal extent (from approximately +17.0 to –2.0mm from the interneural line). Affected regions included the injection sites and extended to nucleus accumbens, claustrum, ventral pallidum, substantia nigra, anterior cingulate cortex, amygdala, entorhinal cortex, and hippocampal formation, as well as prefrontal, orbitofrontal and lateral frontal cortices (areas 8b, 8aD, 46, 47, 11, 13 and 9). The spatial pattern of pathology was heterogeneous: inclusion density varied markedly across regions, with particularly prominent immunoreactivity within the basal ganglia and associated limbic structures, suggestive of region-specific vulnerability.

These patterns are illustrated in Figure 1 across three representative rostrocaudal levels selected to capture the injection site and the anterior and posterior extent of pathological spread; comprehensive coverage across eight rostrocaudal levels and qualitative comparison across timepoints are provided in Supplementary Figures 2–6. No pS129 immunoreactivity was detected in phosphate-buffered saline (PBS)-injected control marmosets across any region or timepoint (see Supplementary Figure 7), confirming that the observed pathology is specific to PFF-induced seeding.

**Figure 1.**
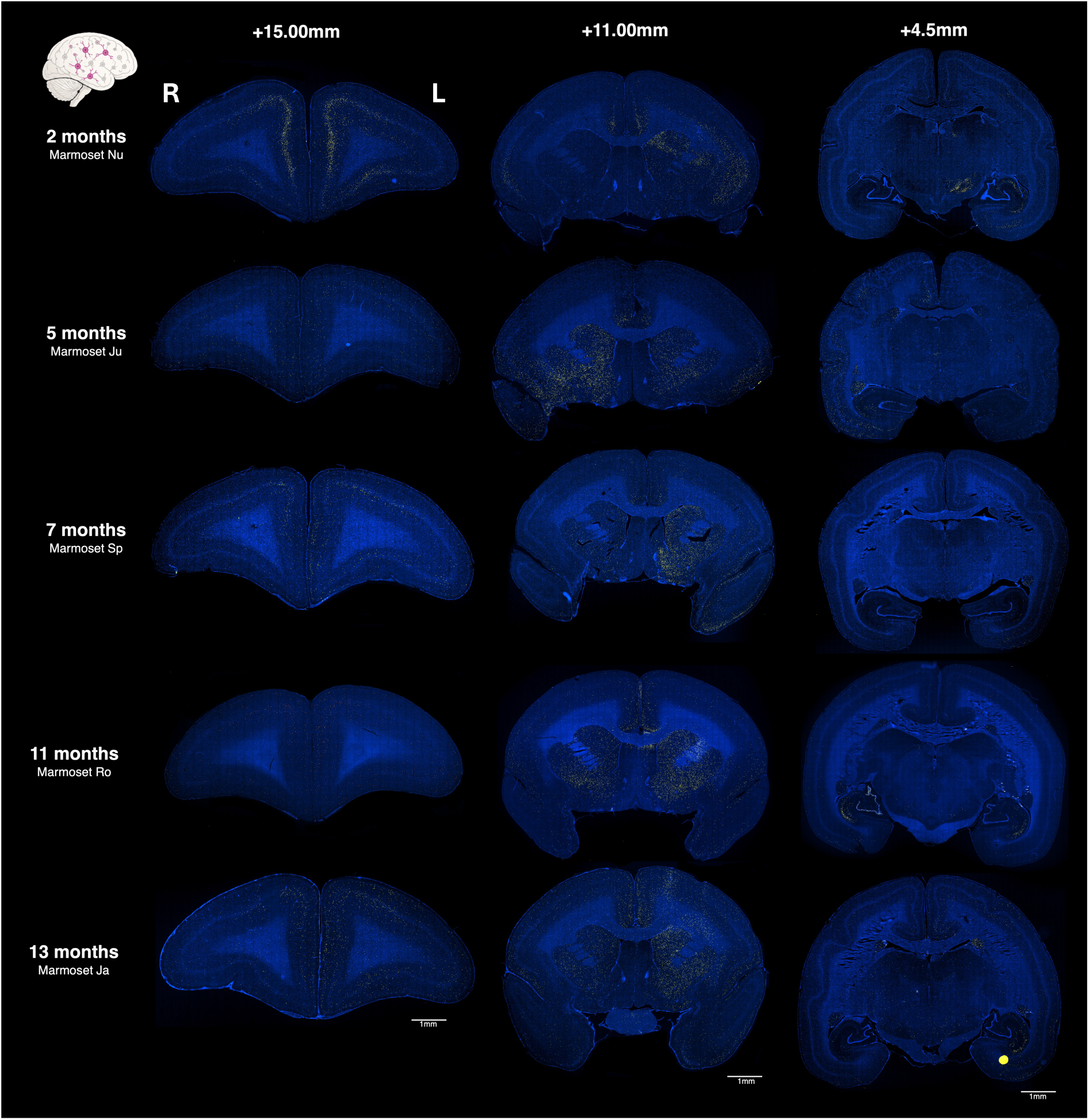
Distribution of pS129-positive αSyn pathology across coronal brain sections from PFF-injected marmosets at multiple post-injection timepoints. Pathological inclusions were detected by immunolabelling against pS129 (magenta); nuclei were counterstained with Hoechst (blue). For each animal, three rostrocaudal levels are shown (+15.0, +11.0, and +4.5 mm from the interaural line), as defined in the Paxinos marmoset brain atlas^18^. Scale bars: 1mm.

High-magnification imaging (63x) provided detailed characterization of pS129-positive inclusion morphology across key regions. Across all areas examined, immunolabelling revealed inclusion-like structures with distinct morphologies, including punctate aggregates, neuritic-like accumulations, and Lewy-body like inclusions. These patterns are illustrated in Figure 2 for four representative regions in the hemisphere ipsilateral to the injection: caudate, putamen, substantia nigra, and hippocampus (CA1).

**Figure 2.**
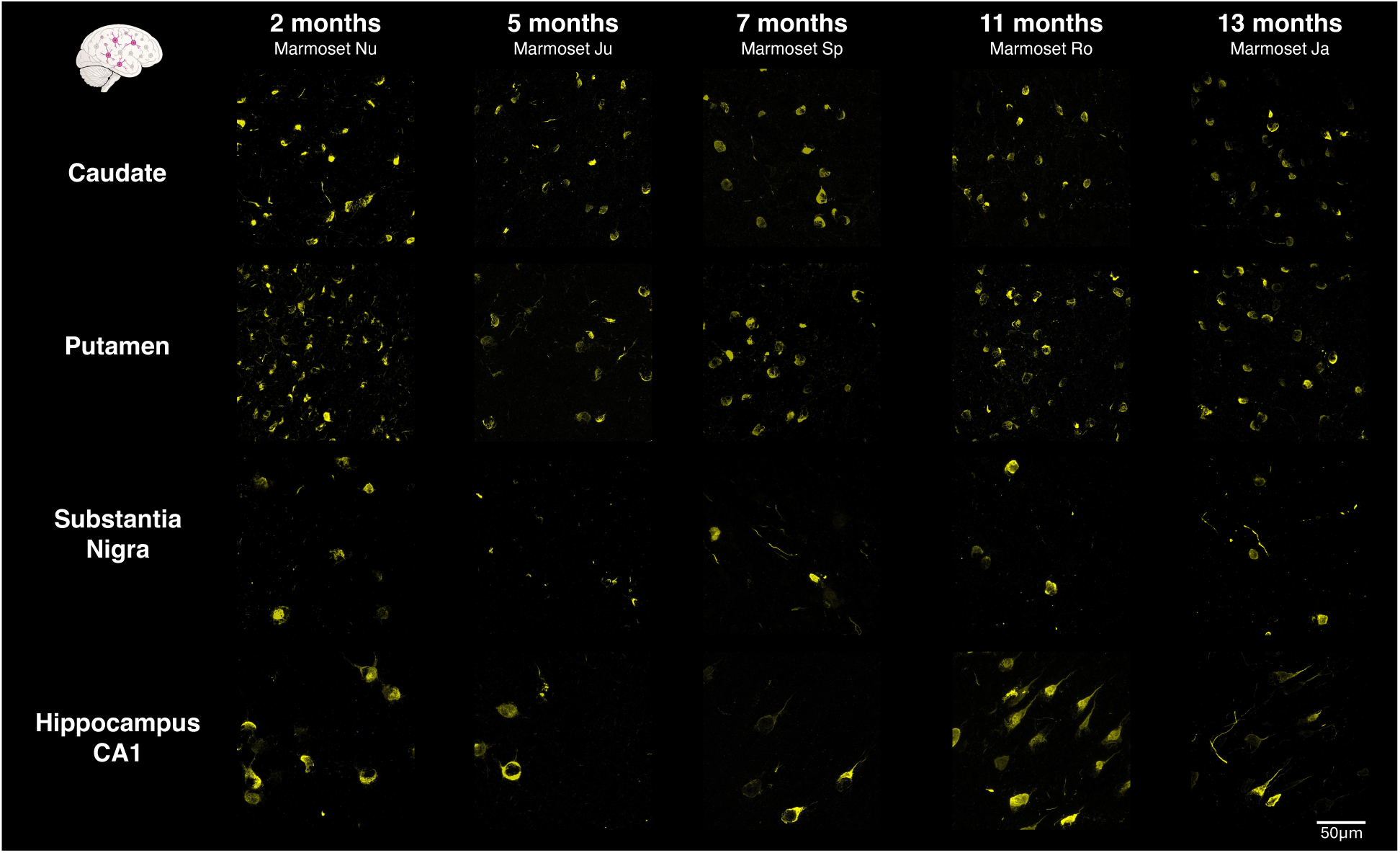
Characterization of pS129 αSyn inclusions in region-specific brain areas. High magnification (63x) immunostaining images showing pS129-positive αSyn (yellow) pathology in selected brain regions, caudate, putamen, substantia nigra and hippocampus, across all time points (2, 5, 7, 11, and 13 months post-injection). Scale bar: 50µm.

In the caudate and putamen, inclusions were abundant from the earliest timepoint examined, consistent with their role as primary injection sites. Early stages were dominated by punctate aggregates, while later timepoints showed a progressive shift toward larger, more morphologically complex Lewy body–like inclusions, suggesting a maturation of pathological deposits over time. In the hippocampus (CA1), pathology evolved from sparse early immunoreactivity to progressive engagement of axonal processes at later timepoints, a pattern consistent with the involvement of long-range projections connecting hippocampal circuits to the broader limbic system. The substantia nigra displayed the most variable pattern across animals and timepoints: inclusions were observed in neuronal cell bodies but also extended to dendritic processes, reflecting a more heterogeneous and potentially more dynamic engagement of nigral neurons compared to striatal structures.

Across all regions, both the abundance and morphological complexity of inclusions increased over time, consistent with progressive accumulation of pathological αSyn. Regional differences in inclusion density and morphology were apparent, with the basal ganglia showing particularly prominent immunoreactivity from early stages, while limbic and nigral involvement became more pronounced at later timepoints.

To assess the integrity of dopaminergic neurons in the substantia nigra across the course of the study, tyrosine hydroxylase (TH) co-immunolabelling was performed in PFF-injected marmosets at all five post-injection timepoints (2, 5, 7, 11, and 13 months; Supplementary Figure 8). At 2 months post-injection, pS129-positive inclusions showed prominent co-localization with TH-positive neuronal cell bodies, indicating early localization of dopaminergic neurons by αSyn pathology. Across subsequent timepoints, the degree of somatic co-localization progressively decreased, accompanied by a relative reduction in pS129 immunoreactivity within cell bodies and a shift toward greater inclusion burden in neuritic processes. Despite this evolving pattern of pathological involvement, TH-positive neurons remained abundant throughout the observation period, including at the latest timepoint examined (13 months post-injection), indicating that dopaminergic neurons were not substantially depleted over the course of the study.

### Deformation-based Morphometry (DBM)

DBM identified widespread structural changes associated with PFF injection, revealed by a significant Group*Timepoint interaction reflecting a greater longitudinal decrease in log-Jacobian values in PFF compared to PBS animals between baseline and 11 months.

The largest cluster was located in the hemisphere ipsilateral to the injection and encompassed key components of the basal ganglia, including the striatum (caudate and putamen), globus pallidus, nucleus accumbens, and extending into the claustrum. Additional clusters were observed in the contralateral caudate and posterior putamen, indicating bilateral involvement of striatal structures.

Beyond the basal ganglia, smaller but consistent clusters were detected in limbic and associative regions, including the anterior cingulate cortex (areas 25, 24a, 24b), ipsilateral orbitofrontal cortex (areas 11 and 13), amygdala, entorhinal cortex, and hippocampal formation. Bilateral effects were also observed in the substantia nigra, alongside more limited clusters in the cerebellum.

These clusters are illustrated in Figure 3A across coronal sections at different interaural levels. Inspection of region-wise trajectories (Figure 3B) revealed that the interaction effects were driven by a progressive decrease in log-Jacobian values in PFF animals, consistent with sustained tissue contraction over time, whereas PBS animals showed relatively stable profiles.

**Figure 3.**
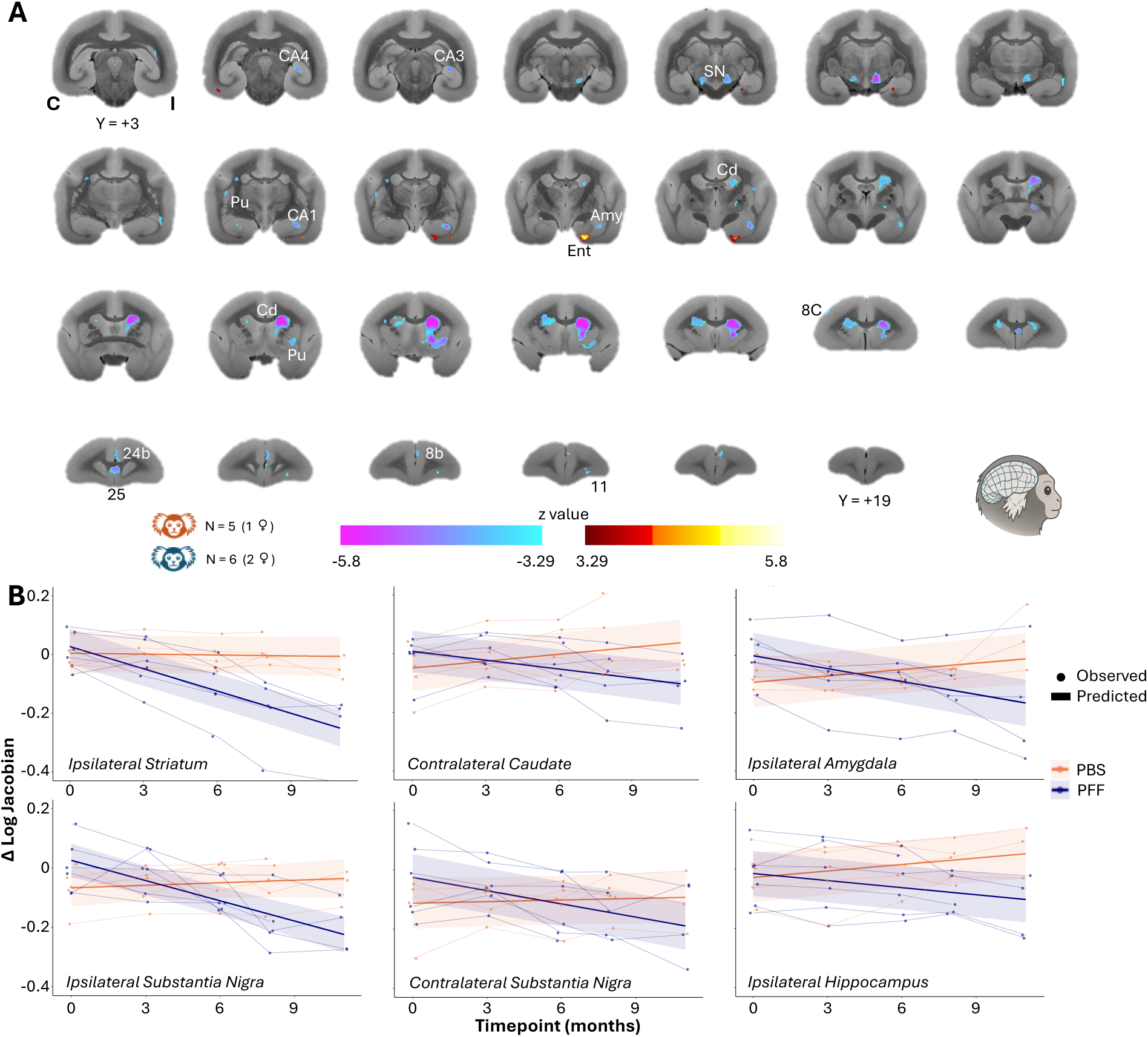
DBM reveals progressive structural divergence between PFF and PBS animals. (A) Coronal sections of the NIH marmoset brain template^19^ (+3 to +19 mm from the interaural line, as defined in the Paxinos marmoset brain atlas^18^) displaying voxelwise results of the Group*Timepoint interaction on log-Jacobian values. Significant clusters indicate regions where the rate of volumetric change over time differs between PFF and PBS animals. Cold colors indicate a more positive slope in PBS animals (relative volume increase or lesser contraction); warm colors indicate a more positive slope in PFF animals. Statistical maps were thresholded at p<0.001 (voxelwise) with a minimum cluster extent of 50 contiguous voxels. Abbreviations: C, contralateral (to the injection site); I, ipsilateral; Cd, caudate; Pu, putamen; CA, cornu ammonis (hippocampus); Amy, amygdala; SN, substantia nigra; Ent, entorhinal cortex. (B) Region-of-interest line plots show log-Jacobian values across timepoints for PBS (orange) and PFF (blue) animals. Solid lines represent group-level trajectories estimated from the linear mixed-effects model, with shaded regions representing the confidence interval; dots represent individual observed values. A negative slope indicates progressive tissue contraction; a positive slope indicates relative expansion.

### Functional Connectivity (FC)

Resting-state FC was assessed using pairwise correlations between the BOLD time series of regions of interest (ROIs) defined according to the Paxinos marmoset cortical atlas, supplemented by subcortical structures; correlation coefficients were Fisher z-transformed for statistical comparisons. To ensure a common reference frame across animals, all analyses are described relative to the injection site (ipsilateral and contralateral) rather than anatomical left and right. This approach yielded group-level FC matrices at three timepoints for each experimental group: baseline, early window (3–6 months post-injection), and late window (8–11 months post-injection); full methodological details are provided in the Methods section. Baseline (∼1 month post-injection) FC matrices revealed pre-existing differences between groups (see Supplementary Information and Supplementary Figure 9). In particular, PFF animals showed stronger FC of auditory and temporal regions in the hemisphere ipsilateral to the injection, both within and across hemispheres, whereas PBS animals exhibited stronger FC within contralateral frontal regions and between these regions and the rest of the brain.

Longitudinal changes in FC (Figure 4) revealed markedly divergent trajectories between groups. PBS animals displayed a generalized increase in FC strength over time, more evident in the hemisphere contralateral to the injection and not restricted to a specific network. In contrast, PFF animals showed a progressive and widespread decline in FC strength, which was more pronounced in the late window (8-11 months post-injection) compared to the early one (3-6 months). Within this global decrease, two partially distinct frontal and prefrontal clusters showed particularly marked reductions in FC. First, a cluster of lateral prefrontal and premotor regions (including areas 8, 46, 47, 9, and portions of area 6) exhibited reduced FC most prominently in the ipsilateral hemisphere, affecting both within-cluster connectivity and interactions with the rest of the brain. Second, orbitofrontal regions showed a progressive and spatially specific decline: areas 10, 14R, and 14C displayed reduced connectivity both within the frontal lobe and towards parietal and occipital areas bilaterally, while areas 11 and 13M showed decreased connectivity within the frontal lobe and with auditory regions, predominantly ipsilaterally but extending contralaterally at the latest timepoint. Notably, areas 11, 13b, and 14 also exhibited reduced connectivity with the ipsilateral and contralateral putamen and thalamus, with the extent of thalamo-frontal disconnection varying across orbitofrontal subregions. Together, these findings indicate a progressive disruption of both lateral prefrontal and orbitofrontal networks, with the latter showing specific involvement of frontostriatal and thalamo-frontal circuits. In contrast, a set of parietal (e.g., PE, PG, PF) and occipital regions showed increased FC in the late window, primarily within the cluster and across hemispheres.

**Figure 4.**
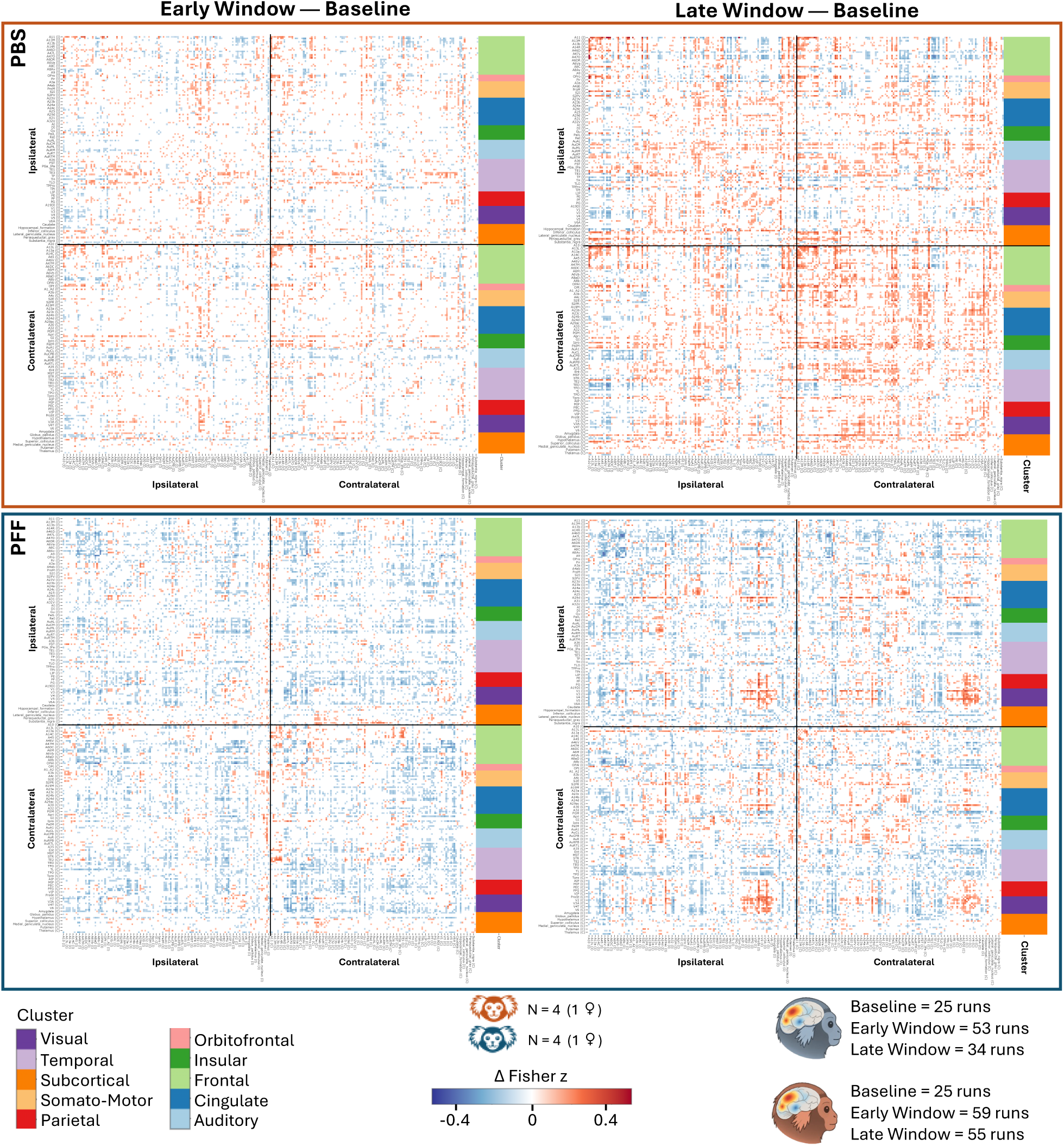
Functional connectivity (FC) matrices reveal divergent longitudinal trajectories in PBS and PFF animals. Delta connectivity matrices (timepoint minus baseline, in Fisher’s z units) are shown for the PBS (top panel, orange frame) and PFF (bottom panel, blue frame) groups. Left panels show the early window (3–6 months post-injection); right panels show the late window (8–11 months post-injection). Blue colors indicate decreases in FC strength relative to baseline; red colors indicate increases. Each matrix is organized into four quadrants reflecting ipsilateral (top left), contralateral (bottom right), and interhemispheric (top right and bottom left, mirrored) connectivity. Colored bars along the matrix Y axis denote anatomical clusters (e.g., light green, frontal regions; pink, orbitofrontal regions) to aid visual interpretation. Regions of interest were drawn from the Paxinos parcellation^18^ of the NIH marmoset brain atlas^19^, supplemented with subcortical structures from the Subcortical Atlas of the Marmoset^20^. For clarity, odd and even region labels are displayed on the x- and y-axes, respectively.

Complementary analyses of global FC changes using summary metrics (mean signed change and Frobenius distance) confirmed that these effects primarily reflected differences in the direction, rather than the magnitude, of connectivity changes between groups (see Supplementary Information, Supplementary Figures 10 and 11, and Supplementary Table 1 for detailed description of these results).

To assess potential confounding effects, temporal signal-to-noise ratio (tSNR) and head motion were computed across timepoints and groups (Supplementary Figure 12). Head motion was quantified using framewise displacement (FD), which summarizes volume-to-volume changes across the six rigid-body motion parameters. No significant effects were observed for tSNR (all effects showing p>0.05), either between groups or across timepoints. In contrast, FD analysis revealed a significant Group*Timepoint interaction, driven by an increase in motion over time within the PBS group, with both early (mean=0.128 mm, p=0.007) and late windows (mean=0.132 mm, p=0.002) showing higher displacement compared to baseline (mean=0.109 mm). No other comparisons were significant. Although volumes with excessive motion were censored during preprocessing, this residual increase in motion may partially contribute to the apparent FC increases observed in PBS animals. However, the absence of tSNR differences and the spatial specificity of the FC patterns suggest that the main findings are unlikely to be solely explained by motion-related artifacts.

Seed-based analyses (Figure 5) further characterized the spatial specificity of connectivity changes by examining four ipsilateral seeds: mediodorsal (MD) thalamus, caudate, hippocampal formation, and a frontal cluster encompassing areas 46 and 8, the regions showing the strongest decrease in FC in the PFF sample.

**Figure 5.**
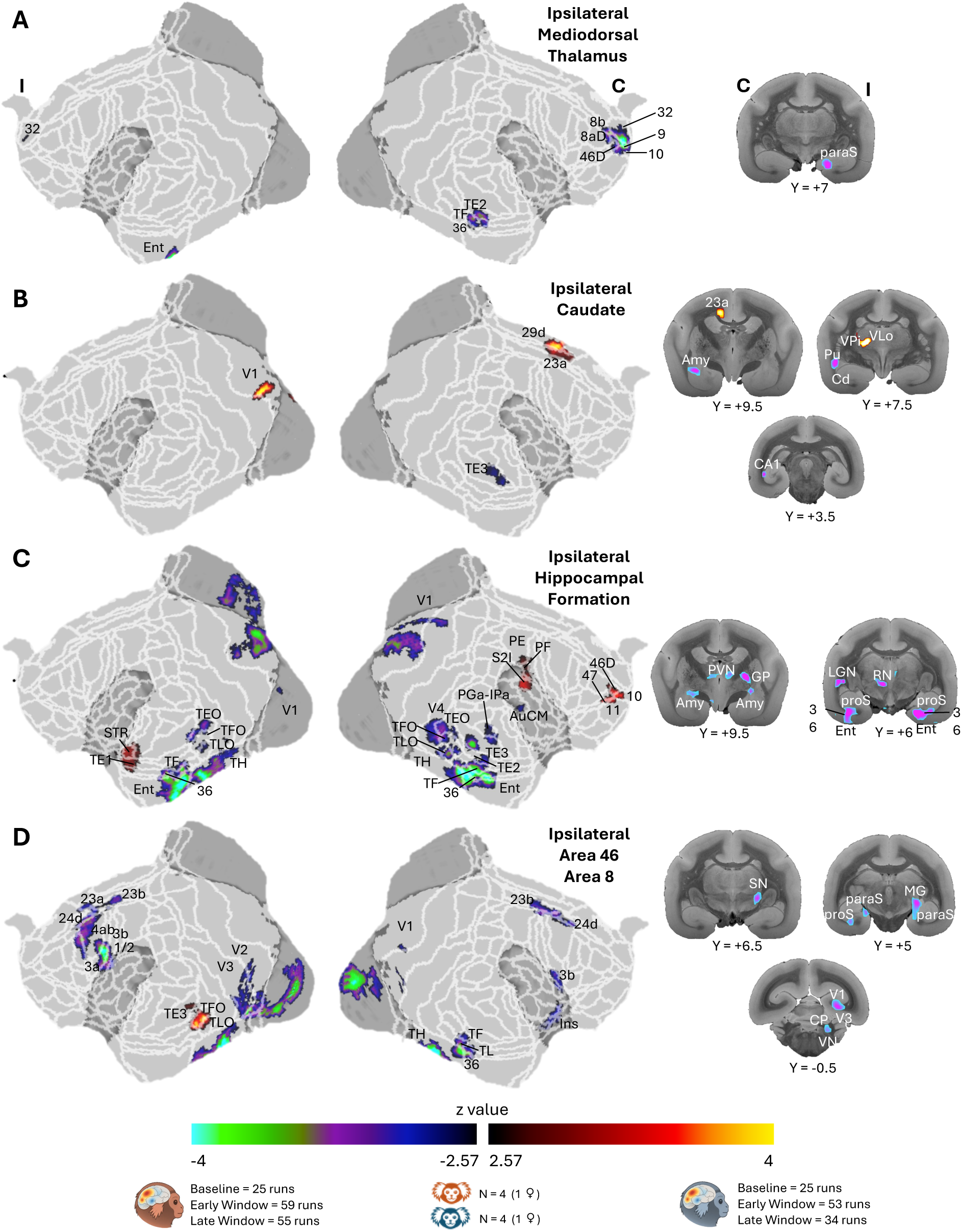
Seed-based analyses reveal network-specific changes in functional connectivity in PFF animals. Voxelwise maps show the difference in longitudinal slopes (PFF minus PBS) for functional connectivity seeded from four ipsilateral regions of interest: (A) mediodorsal thalamus, (B) caudate, (C) hippocampal formation, and (D) a frontal cluster encompassing areas 8 and 46. Significant clusters of the Group*Timepoint interaction are thresholded at p<0.01 (voxelwise) with cluster-level correction at a<0.05 and a minimum cluster extent of 50 voxels. Cold colors indicate a more negative slope in PFF animals (greater decrease in functional connectivity over time); warm colors indicate a more positive slope in PFF animals (relative increase). Results are displayed on cortical flat surfaces and on selected coronal sections of the NIH marmoset brain template^19^. White lines on flat maps delineate the Paxinos parcellation^18^. Abbreviations: I, ipsilateral; C, contralateral; Cd, caudate; Pu, putamen; Amy, amygdala; Ent, entorhinal cortex; paraS, parasubiculum; proS, prosubiculum; Ins, insula; GP, globus pallidus; SN, substantia nigra; RN, red nucleus; LGN, lateral geniculate nucleus; MG, medial geniculate nucleus; PVN, paraventricular nucleus; VP/VL, ventral posterior and ventral lateral thalamic nuclei.

For the MD thalamus, the Group*Timepoint interaction revealed reduced connectivity in PFF animals with contralateral prefrontal regions (areas 8b, 8aD, 9, 10, 11, 32 and 46D), as well as with temporal areas (TE2, TF and caudal area 36). Additional reductions were observed subcortically in the presubiculum and parasubiculum, and in the cerebellum, particularly within contralateral Crus I and II.

The ipsilateral caudate seed showed a mixed pattern of connectivity changes in PFF animals. Increased connectivity was observed with posterior cingulate regions (areas 23a, 29d and 30), a small occipital cluster in V1, and contralateral thalamic nuclei. In contrast, decreased connectivity was found with the contralateral putamen, amygdala, caudate, and temporal area TE3.

The hippocampal formation exhibited the most extensive pattern of altered connectivity, characterized predominantly by widespread reductions in PFF animals. These encompassed multiple subcortical regions, including globus pallidus, basal forebrain cholinergic nuclei, amygdala, hypothalamus, thalamic nuclei (including MD) and bilateral hippocampal structures, as well as contralateral caudate and midbrain regions. Reductions extended to the cerebellum and to widespread cortical territories across occipital, temporal and parietal lobes. More limited increases in connectivity were observed in contralateral parietal (S2I, PE, PF) and prefrontal (10, 11, 46D, 47) regions and in ipsilateral temporal cortex (STR, TE1).

Finally, the frontal seed (including areas 46 and 8) revealed a broad pattern of reduced connectivity in PFF animals, involving cingulate (23a, 23b, 24d), somato-motor (1/2, 3a, 3b and 4ab), insular, temporal (TF, TL, TH, 36) and visual cortices (V1, V2, V3), as well as subcortical regions including the medial geniculate nucleus, substantia nigra and parasubiculum. A smaller cluster of increased connectivity was observed within ipsilateral temporal areas (TFO, TLO and TE3).

### Regional Homogeneity (ReHo)

ReHo measures the local synchrony of BOLD signal fluctuations within neighboring voxels, providing an index of local circuit coordination independent of long-range connectivity^21^. ReHo analyses revealed a consistent pattern of reduced local functional synchronization in PFF animals compared to PBS (Figure 6A). Interaction maps (timepoint PFF – baseline PFF vs timepoint PBS – baseline PBS) computed at the early and late windows showed largely overlapping spatial patterns, with the extent and magnitude of effects increasing in the late interval, indicating a progressive decline in ReHo over time following PFF injection.

**Figure 6.**
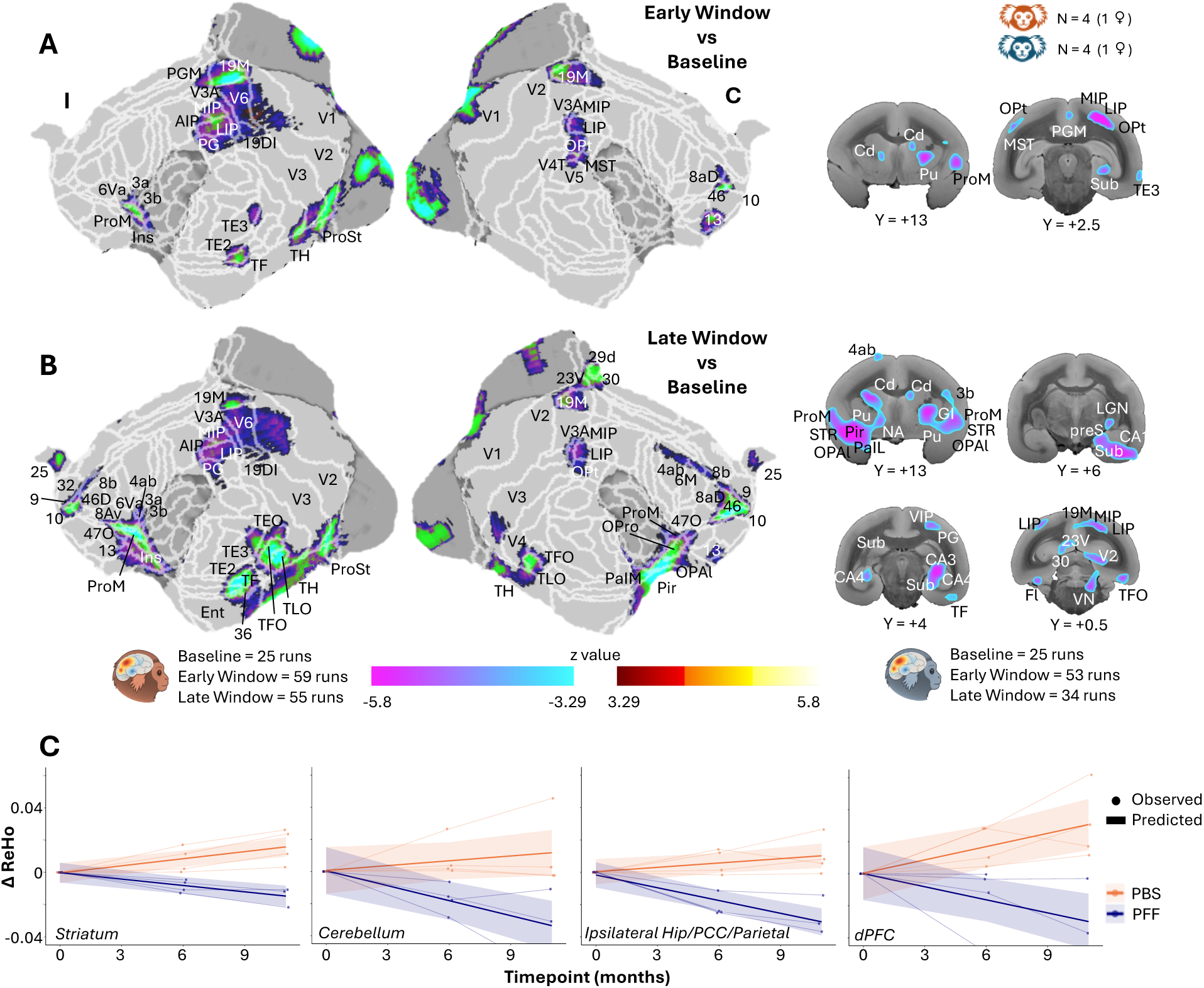
Regional homogeneity analysis reveals widespread reductions in local functional synchrony in PFF animals. Voxelwise maps of the Group*Timepoint interaction in ReHo are shown on cortical flat surfaces and selected coronal sections of the NIH marmoset brain template^19^ with white lines delineating the Paxinos parcellation^18^, for the early (panel A; 3–6 months minus baseline) and late (panel B; 8–11 months minus baseline) windows. Cold colors indicate a more negative ReHo difference in PFF animals relative to PBS animals; warm colors indicate a more positive difference. Statistical maps are thresholded at p<0.001 (voxelwise) with cluster-level correction at α<0.05 and a minimum cluster extent of 50 voxels. Panel C shows delta ReHo trajectories across timepoints for four representative regions of interest — bilateral striatum, bilateral cerebellum, an ipsilateral cluster encompassing hippocampal formation, posterior cingulate cortex and parietal cortex, and bilateral dorsal prefrontal cortex (left to right) — for PFF (blue) and PBS (orange) animals. Solid lines represent group-level trajectories estimated from the linear mixed-effects model; dots represent individual observed values. A negative slope indicates a decrease in local synchrony over time. Abbreviations: I, ipsilateral; C, contralateral; Cd, caudate; Pu, putamen; NA, nucleus accumbens; Sub, subiculum; Pir, piriform cortex; Ins, insula; VN, vestibular nucleus.

These reductions encompassed widespread cortical territories, including frontal regions (including areas 6Va, 8, 9, 10, 25, 32 and 46), somato-motor cortex (areas 4ab, 3a and 3b), parietal regions (including MIP, LIP, AIP and PG), temporal cortex (including TE2, TE3, TEO, TF and TFO), visual areas (V1, V2, V3, ProSt and 19M), and cingulate regions (areas 29d, 30 and 23V).

Subcortically, significant reductions in ReHo were observed in the bilateral caudate and putamen, the lateral geniculate nucleus (LGN), and the hippocampal formation. Additional decreases were identified in the ipsilateral cerebellum, particularly within the paramedian lobule.

To illustrate the temporal dynamics underlying these effects, representative clusters were extracted and plotted (Figure 6B). Across all regions, the significant Group*Timepoint interactions were driven by a marked decrease in ReHo in PFF animals, whereas PBS animals showed stable or slightly increasing trajectories over time. This pattern was consistently observed in clusters encompassing the striatum, dorsal prefrontal cortex, cerebellum, and a broader network including hippocampal, posterior cingulate and parietal regions.

### Actimetry

Motor activity and sleep were monitored longitudinally using a veterinary actigraph attached to a collar (McGill University) or headpost-mounted holder (Western University), with recordings collected over weekends every two months starting one month post-injection. Raw activity data were classified into sleep and wake states using an activity threshold approach, and a set of night-time and daytime metrics were derived for each session; full methodological details are provided in the Methods section. Longitudinal actimetry analyses did not reveal significant effects of Group, Timepoint, or their interaction for any of the examined metrics, including night-time measures (total sleep duration, sleep latency, transitions) and daytime measures (mean activity, number of naps, nap duration) (Figure 7). For all metrics, linear mixed-effects models with Group and Timepoint as fixed effects and subject as a random intercept yielded no significant main effects or interactions (all p>0.05), and no consistent or coherent pattern of differences emerged between PFF and PBS animals across timepoints. Variability across timepoints appeared irregular and likely reflects the limited sample size rather than systematic group-level effects. Consistent with these findings, qualitative observations throughout the study period did not reveal obvious motor abnormalities such as tremor, bradykinesia, or other behavioural disturbances even at the latest timepoint examined (13 months post-injection).

**Figure 7.**
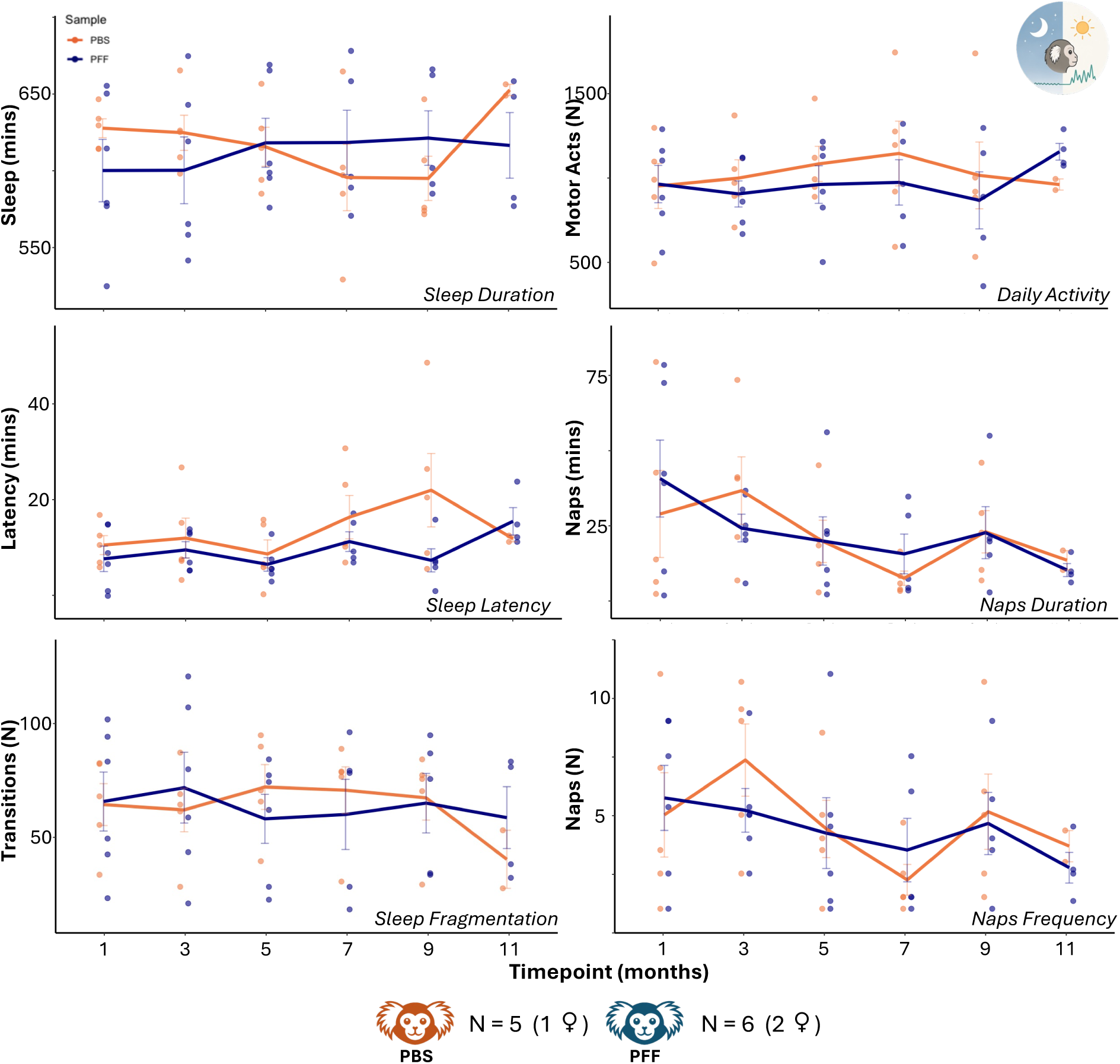
No significant group differences in sleep or motor activity across the study period. Group-level mean trajectories (solid lines) and individual values (colored dots) are shown for PFF (blue) and PBS (orange) animals across six actimetry-derived measures acquired over three nights and two days (from Friday evening to Monday morning). Left column (top to bottom): sleep duration (minutes), sleep latency (minutes), and sleep fragmentation (number of sleep/wake transitions per night). Right column (top to bottom): daily motor activity (number of actimetry-detected motor events during the day), mean nap duration (minutes), and nap frequency (number of daytime naps). No significant differences between groups were detected for any measure. Colored bars represent the standard error of the mean. Note: night-time was defined as 19:00–07:00 h at Western University and 20:00–08:00 h at McGill University.

### Pairwise Visual Discrimination (PVD) Task

Impaired cognitive flexibility is a recognized feature of synucleinopathies^22,23^, and reversal learning paradigms offer a well-validated and cross-species approach to its assessment^24,25^. Therefore, cognitive performance in the touchscreen-based PVD task was assessed longitudinally to determine whether PFF injection affected learning and cognitive flexibility (Figure 8). Performance did not differ between groups during the learning phase, with both PBS and PFF animals acquiring the stimulus–reward association at comparable rates across all timepoints. On average, monkeys reached criterion within fewer than 200 trials, indicating preserved learning ability in the PFF group. In contrast, the reversal phase revealed a significant Group*Timepoint interaction (F_(4, 20.49)_ = 3.76, p = 0.019). Post hoc comparisons confirmed that PBS and PFF animals performed similarly at baseline (PBS=309±65 (mean±SEM), PFF=229±19, p=0.26), but differed at 5 (PBS=195±30.8, PFF=352±63, p=0.035), 7 (PBS=139±32.1, PFF=302±46.6, p=0.047), and 9 months (PBS=167±29.1, PFF=332±21.5, p=0.043), with PBS animals requiring fewer trials to reach criterion (90% correct responses over the last 50 trials performed). A similar pattern was observed at 11 months (PBS=180, PFF=331.5±21.5), although this comparison did not reach significance (p=0.50) and should be interpreted cautiously and as purely exploratory due to the limited sample size (n = 2 PFF vs 1 PBS). No significant effects were observed for error rates (both for regressive and perseverative errors) or task engagement (number of trials performed in the first 3 minutes of each session) in either learning or reversal phases (Supplementary Figure 13).

**Figure 8.**
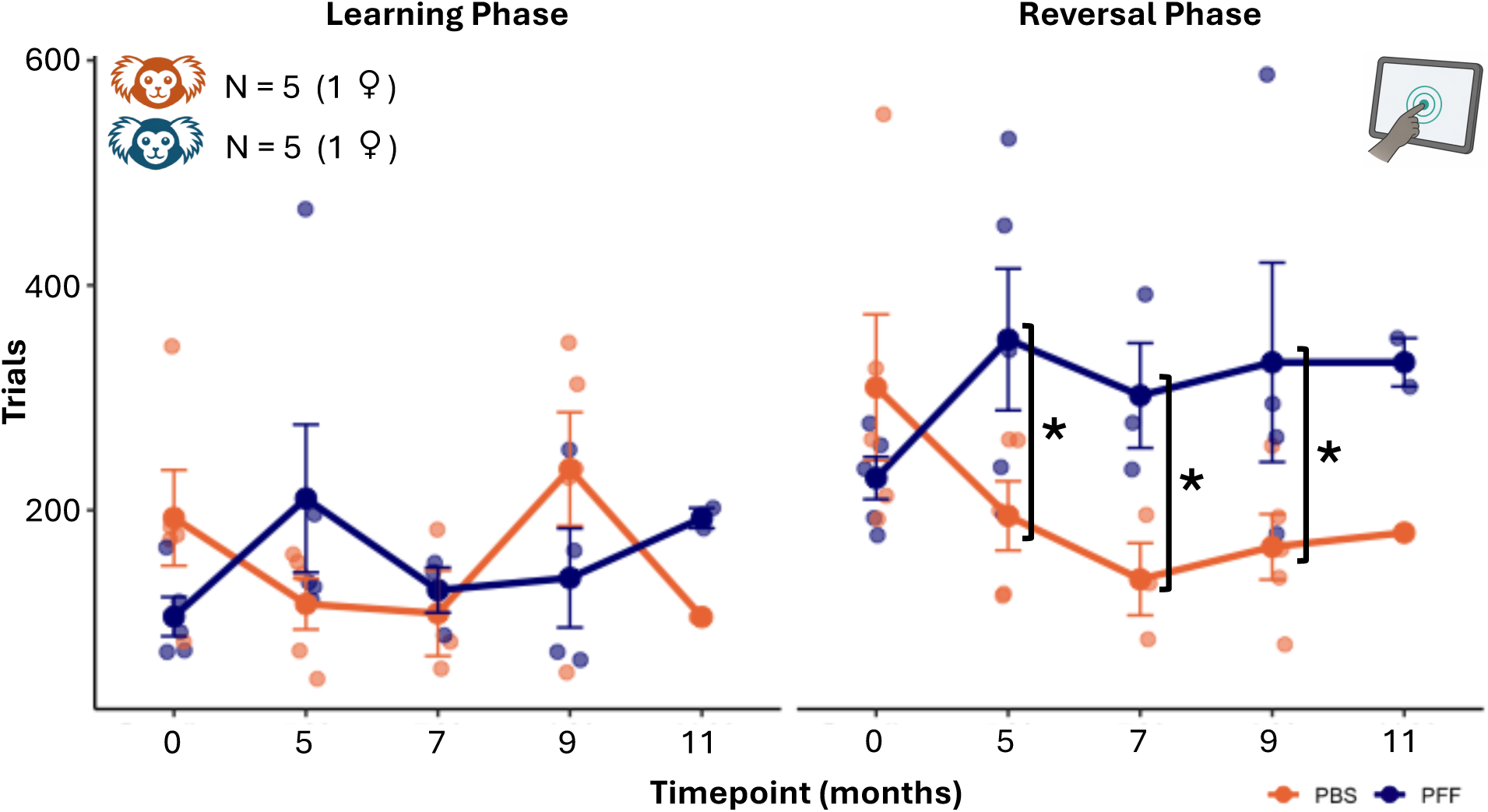
Selective impairment in reversal learning in PFF animals. Performance on the touchscreen-based PVD task is shown for PBS (orange) and PFF (blue) animals across timepoints, expressed as the number of trials required to reach criterion (90% correct responses over the last 50 trials). Left panel: learning phase, in which one stimulus (S+) was consistently rewarded and the other (S−) was not. Right panel: reversal phase, in which the stimulus-reward contingencies were reversed. Solid lines represent group means; dots represent individual values; error bars represent the standard error of the mean. Statistical comparisons were performed using separate linear mixed-effects models for the learning and reversal phases; significant effects were followed by post-hoc comparisons corrected with the Holm-Bonferroni method. Asterisks denote group differences at p<0.05. Note: animal numbers per timepoint vary due to planned removal for histological assessment; the final timepoint is exploratory only (PFF, n=2; PBS, n=1).

## DISCUSSION

Using a longitudinal, multimodal framework combining histology, structural MRI, resting-state functional connectivity, regional synchrony measures, and behavioural assessments, we characterized the progressive consequences of striatal αSyn seeding in the common marmoset. Across all modalities, the results converge on a coherent picture: PFF injection initiates a cascade of pathological change that spreads beyond the injection site, driving distributed structural and functional reorganization and culminating in cognitive impairment without obvious motor deficits. Rather than a focal or purely motor syndrome, these findings support the view that α-synucleinopathy in this model unfolds as a systems-level disorder, with its primary impact on large-scale brain networks supporting executive function and limbic processing^26^.

Histological analyses demonstrated a time-dependent spreading of phospho-S129 αSyn pathology from the caudate and putamen injection sites to widespread cortical and subcortical regions. Propagation was spatially heterogeneous, with preferential involvement of limbic and associative areas including the hippocampus, amygdala, and prefrontal cortex, consistent with a spreading along connectivity-constrained pathways already described in literature^4,9,11,15,27^. The presence of inclusion-like structures with morphological features of Lewy body–type pathology supports the relevance of this model to the broader class of synucleinopathies. In particular, the prominent engagement of hippocampal and prefrontal regions is more consistent with synucleinopathy phenotypes involving early cognitive and limbic dysfunction, such as dementia with Lewy bodies (DLB), than with strictly nigrostriatal-predominant forms of Parkinson’s disease (PD)^5,28–30^. Together, these observations indicate that pathological spread in this model follows a distributed, network-embedded trajectory rather than a strictly focal or dopaminergic one.

Consistent with the histological pattern, deformation-based morphometry revealed a progressive tissue contraction in PFF marmosets. Effects were most pronounced in the ipsilateral striatum and associated nuclei, but extended to contralateral striatum, anterior cingulate, orbitofrontal cortex, hippocampus, amygdala, and substantia nigra — a spatial distribution that closely mirrored pathological spread across the same time window. The congruence between structural atrophy and pathology extent supports the interpretation that volume loss in this model is driven by advancing neurodegeneration rather than non-specific injection effects, and further underscores the network-level nature of the degenerative process. Notably, comparable atrophy patterns have recently been reported in longitudinal PFF mouse models, where structural contraction extended beyond the injection site to frontal, hippocampal, thalamic, and limbic regions^17^, suggesting that the large-scale structural consequences of α-synuclein seeding may follow conserved organizational principles across species. However, whereas previous results also reported bilateral somatomotor cortex atrophy (consistent with the motor deficits commonly observed in rodent PFF models^11,17^), somatomotor regions were not among the affected structures in the present dataset. This relative sparing is consistent with the absence of overt motor symptoms in our animals even at the latest timepoints examined, and further supports the interpretation that the present model captures a predominantly cognitive and limbic stage of synucleinopathy, distinct from the motor-dominant profile typically associated with classical PD^5,28–30^.

Although qualitative inspection of baseline connectivity matrices suggested pre-existing group differences, formal longitudinal analyses modelled within-subject change and are therefore unaffected by this observation. Longitudinally, PFF and PBS animals diverged markedly: control marmosets showed a modest increase in connectivity over time, which may partly reflect methodological factors such as motion, though this was not formally tested. By contrast, PFF marmosets exhibited a progressive and widespread decline in connectivity, particularly within frontal and prefrontal regions (areas 8, 9, 46, and 47). Notably, posterior parietal and occipital regions showed relative increases in connectivity at later timepoints, suggesting a reorganization of large-scale network topology rather than a uniform loss of functional coupling. Similar patterns of frontal disconnection accompanied by relative preservation or enhancement of posterior connectivity have been reported in synucleinopathies and other neurodegenerative disorders and may reflect compensatory or maladaptive redistribution of network dynamics^31–33^. These results are unlikely to reflect acquisition artefacts, given that all functional imaging data were collected at a single site using a consistent protocol and no group or timepoint differences in tSNR were observed.

Seed-based analyses confirmed that these connectivity changes reflected disruption of specific large-scale circuits. Reduced connectivity of the mediodorsal thalamus with prefrontal and temporal regions, widespread hippocampal disconnection, and broad decreases seeded from frontal areas 8 and 46 collectively point to a progressive breakdown of fronto-striato-thalamo-cortical and limbic networks. These circuits are central to executive function and cognitive flexibility^34–36^, and their disruption constitutes a plausible network-level substrate for the behavioral alterations described below^37^. The concurrent increases in posterior connectivity may reflect compensatory plasticity, disinhibition following frontal network breakdown, or a redistribution of network dynamics^31–33^, though the mechanistic basis of this divergent pattern remains to be established and would require a more targeted follow-up.

Regional homogeneity (ReHo) analyses provided complementary evidence for progressive functional deterioration at the level of local circuit synchrony. PFF marmosets showed widespread reductions in ReHo across frontal, parietal, temporal, and visual cortices, as well as subcortical structures including the striatum, hippocampus, and lateral geniculate nucleus; PBS animals remained stable throughout. Although ReHo abnormalities in Parkinson’s disease and dementia with Lewy bodies are often reported within sensorimotor networks^38–40^, the present pattern was characterized more prominently by fronto-striatal and limbic involvement, consistent with the broader cognitive and associative phenotype observed in this model. Notably, the spatial extent and magnitude of ReHo reductions increased from the early to the late interval, paralleling reports of progressive local synchrony deterioration across disease stages in human synucleinopathies^41^. Critically, ReHo reductions were detectable at earlier timepoints than some long-range connectivity effects and intensified over the course of the study, suggesting that local circuit dysfunction may precede (or at least accompany) the emergence of large-scale network disruption. This temporal pattern is consistent with a hierarchical model in which αSyn pathology first perturbs local neuronal coordination before propagating to impair long-range communication between regions.

Despite extensive structural and functional alterations, neither actimetry nor observational assessments revealed overt motor deficits or sleep disturbances in PFF animals. Longitudinal TH immunolabelling in the substantia nigra of PFF-injected animals across all timepoints (2–13 months post-injection) revealed persistently abundant dopaminergic neurons throughout the observation period, despite the progressive presence of pS129-positive inclusions in both cell bodies and neuritic processes (see Results). The preservation of nigral TH immunoreactivity across timepoints is consistent with the absence of motor symptoms and suggests that αSyn-driven pathology has not yet reached the threshold of dopaminergic loss typically required for overt motor symptom emergence in synucleinopathies. Notably, the longitudinal pattern also revealed a progressive reduction in somatic pS129 immunoreactivity within TH-positive neurons, accompanied by a relative shift of inclusion burden toward neuritic processes. While highly speculative, and based on a single animal per timepoint, precluding firm conclusions, this evolving pattern may suggest a partial resolution or redistribution of nigral pathology over time, which could contribute to the sustained preservation of dopaminergic neurons and the absence of motor symptoms. This hypothesis warrants direct investigation in future studies with larger cohorts and extended follow-up. The absence of sleep disturbances in PFF animals similarly points to an early or incomplete stage of pathological progression. Sleep disturbances, particularly REM sleep behavior disorder (RBD), are recognized as early clinical manifestations of PD and DLB and are thought to reflect early involvement of brainstem REM-regulating circuits^42,43^. The absence of sleep phenotypes in the present dataset is therefore consistent with the apparent lack of prominent brainstem involvement at this stage of the model. However, more subtle alterations in sleep architecture cannot be excluded, given the limited sensitivity of actimetry to detect them. This interpretation is broadly consistent with prior PFF-based animal studies, in which sleep phenotypes are variable and appear most robust in models involving prominent brainstem REM-regulating nuclei^44,45^.

In contrast to the preserved motor profile, cognitive testing revealed a selective and reproducible impairment in reversal learning in PFF animals, emerging in the absence of changes in initial discrimination learning, error rates, or task engagement. This specificity argues against a generalized performance decrement and instead indicates a targeted disruption of adaptive, flexibility-dependent behavior. The reversal learning deficit is consistent with dysfunction of prefrontal and fronto-striatal circuits^34–37^, which were among the most affected networks in both functional connectivity and ReHo analyses. In particular, altered connectivity of areas 8 and 46 and the mediodorsal thalamus aligns with established substrates of behavioral flexibility and executive control in primates^37,46–48^.

Complementing this fronto-striatal account, histological and functional evidence also implicates the orbitofrontal cortex (OFC) as a contributor to the observed cognitive deficit. pS129-positive inclusions were detected in orbitofrontal areas 11 and 13, accompanied by progressive structural atrophy in these same regions. Functional connectivity analyses further revealed a longitudinal decline in connectivity involving OFC areas 10, 11, 13, and 14 in PFF animals. The OFC is a well-established substrate of reversal learning and cognitive flexibility across rodents^25,49^, marmosets^50^, and humans^51,52^, where it is thought to support the updating of stimulus-reward associations — precisely the computation disrupted in the reversal phase of the pairwise visual discrimination task. The convergence of inclusion pathology, structural atrophy, and functional disconnection within OFC therefore provides an additional and anatomically specific circuit-level account of the reversal learning impairment, complementary to the broader fronto-striatal and thalamic disruption described above.

The concurrent hippocampal disconnection further suggests that interactions between memory-related and executive systems may also be impaired, potentially compounding the flexibility deficit. Notably, recent touchscreen-based studies in α-synuclein PFF mouse models have similarly reported selective impairments in cognitive flexibility^53^, sometimes emerging prior to overt motor abnormalities^17^, suggesting that executive dysfunction may represent an early and cross-species consequence of network-level synucleinopathy. Within this framework, the absence of clear motor deficits in the present cohort may reflect an earlier stage of pathological progression, in which fronto-limbic and associative circuits (including prefrontal, thalamic and orbitofrontal systems) are functionally compromised before sufficient nigrostriatal degeneration accumulates to produce overt motor symptoms. Taken together, these findings establish a direct link between network-level dysfunction and behavioral outcome, supporting the conclusion that cognitive impairment in this model arises from distributed circuit disruption rather than localized damage.

Integrating these observations, the present results support a model of αSyn–driven pathology in which striatal seeding initiates a progressive, multi-scale cascade of brain reorganization. This cascade, while not fully resolved in its temporal ordering, encompasses: (i) the accumulation and trans-regional spread of pathological inclusions; (ii) regionally specific structural contraction; (iii) disruption and topological reorganization of large-scale functional networks; (iv) breakdown of local functional coherence; and (v) selective cognitive impairment. The relative preservation of motor behavior, despite significant basal ganglia involvement, suggests that different functional domains have distinct vulnerability thresholds. Frontal and limbic networks supporting cognitive flexibility appear highly sensitive to early disruption, as evidenced by convergent functional and behavioral findings, whereas motor systems may remain functionally preserved due to their reliance on distributed, redundant circuitry and the well-established requirement for substantial nigrostriatal dopaminergic loss before overt motor symptoms emerge^54,55^.

From a translational perspective, the prominence of frontal and limbic network disruption, the presence of Lewy body–like inclusions, and the selective cognitive impairment in the absence of overt motor symptoms collectively suggest that the pathological process initiated in this model shares features with synucleinopathies characterized by early cognitive and limbic involvement, such as DLB or prodromal Parkinsonian disorders^5,28–30^. However, this model does not fully recapitulate the clinical or neuropathological profile of any single condition, and — to our knowledge — no established framework yet exists for classifying α-synucleinopathy phenotypes in the common marmoset. Rather than mapping the present findings onto a predefined disease category, we propose that this model opens a tractable path for systematically investigating which aspects of human synucleinopathy are faithfully reproduced, and under what conditions. The longitudinal, multimodal approach employed here provides the methodological foundation for such future work, including the identification of imaging and behavioral markers that may help discriminate between disease-relevant trajectories as the field matures. Moreover, it also establishes a translational primate framework in which the effects of emerging treatments targeting αSyn aggregation and propagation can be evaluated across molecular, circuit, and behavioral scales.

Several limitations warrant consideration. Sample sizes for longitudinal imaging and behavioral analyses were modest, which may have reduced sensitivity to detect subtle effects — particularly for subgroup-level imaging comparisons — and precludes strong conclusions about effect sizes or individual variability. Structural MRI data were acquired across two sites, though preprocessing was standardized. As an induced model, PFF injection may not fully recapitulate the temporal dynamics or spatially graded onset of endogenous synucleinopathy. Differences in actimetry configuration across sites and the relatively limited duration of behavioral follow-up (e.g., ∼1 year in this present study) may also have reduced sensitivity to emerging motor or sleep phenotypes. Importantly, dopaminergic integrity was not directly assessed in the present study — for instance through tyrosine hydroxylase immunohistochemistry or dopamine transporter imaging — and the relative preservation of motor behavior therefore cannot be attributed to a specific degree of nigrostriatal sparing. Future work should incorporate direct measures of dopaminergic function to clarify the relationship between striatal pathology, dopaminergic loss, and motor threshold effects. More broadly, extended longitudinal follow-up with larger cohorts will be needed to determine whether motor deficits emerge at later stages and to establish more precise causal relationships between network dysfunction and behavioral outcomes.

This study demonstrates that αSyn seeding in the marmoset striatum initiates a progressive, multi-scale reorganization of brain structure and function, characterized by widespread network disruption and selective cognitive impairment in the absence of overt motor deficits. The convergence of histological, structural, and functional evidence across modalities establishes this as a systems-level process, with the fronto-limbic axis emerging as particularly vulnerable at this stage of pathological progression. These findings highlight the importance of conceptualizing synucleinopathies as network disorders and underscore the value of longitudinal non-human primate models for bridging molecular pathology with circuit-level and behavioral outcomes.

## METHODS

### Subjects

Sixteen adult common marmosets (*Callithrix jacchus*; 4 females) were included in this study. The mean age at the time of injection was 3.7 years (min=2, max=6.5 years, see Supplementary Table 2 for full details on the sample). Three marmosets (including one female) were tested at McGill University, while the remaining animals were tested at Western University. Animals were housed under standard conditions with comparable environmental enrichment across sites, including branches, ropes, hammocks, shelters, and toys. A 12 h light/dark cycle was maintained at both sites, with lights on from 07:00 to 19:00 at Western University and from 08:00 to 20:00 at McGill University. Animals received intracerebral injections of either phosphate-buffered saline (PBS) or preformed αSyn fibrils (PFF); further details are provided in the following section. Following injection, animals underwent a combination of experimental procedures, including resting-state fMRI, anatomical MRI, touchscreen-based behavioral testing, and continuous actimetry recording (see Supplementary Figure 1 for a detailed overview of the methods and the timepoints employed). A detailed overview of each animal, including injection type and the specific modalities acquired per subject, is provided in Supplementary Table 2. Animals remained under close veterinary supervision throughout the study. All experimental procedures were conducted in accordance with the guidelines of the Canadian Council on Animal Care and were approved by the Animal Care Committees of Western University and McGill University.

### Preparation of recombinant marmoset a-synuclein preformed fibrils (PFF)

Recombinant marmoset αSyn monomers were produced in ClearColi BL21(DE3) *E. coli*. Transformed bacteria carrying the GST-tagged full-length marmoset αSyn plasmid were selected with ampicillin prior to protein expression in LB medium. At an optical density (OD₆₀₀) of 0.6, protein expression was induced with 300 µM IPTG, and cultures were incubated overnight at 16 °C with shaking at 200 rpm. Cells were harvested by centrifugation and resuspended in ice-cold resuspension buffer (25 mM Tris, pH 8.0, 400 mM NaCl, 5% glycerol, 0.5% Triton X-100, protease inhibitors, and 1 mM DTT). Following sonication and clarification by centrifugation, lysates were applied to glutathione Sepharose 4B resin, and bound GST–αSyn was eluted with 20 mM glutathione in wash buffer.

GST tags were removed by overnight cleavage with GST-3C protease (1:50 mass ratio) at 4 °C. The untagged marmoset αSyn was recovered from the flow-through of a GSTrap 4B column and further purified by size-exclusion chromatography using a Superdex 200 16/600 column on an ÄKTA Pure system. Purified protein was exchanged to sterile PBS, pH7.4 (Wisent) and concentrated to 5 mg/mL, filtered through a 0.22 µm membrane, aliquoted (500 µL), and stored at −80 °C.

To generate PFFs, a 500 µL aliquot of monomeric marmoset αSyn (5mg/mL) was sealed with Parafilm and incubated at 37 °C in a digital heating shaking dry bath at 1000 rpm for 7 days. Synuclein aggregates appeared turbid following incubation. Formation of β-sheet-rich amyloid structures and fibrillar morphology was subsequently confirmed by Thioflavin-T assay and transmission electron microscopy (TEM), respectively (Supplementary Figure 14A). The fibrils were sonicated by a Bioruptor Pico (Diagenode) at least 70 cycles of 30 s ON/30 s OFF, 5°C water circulation. Resulting PFFs were analyzed by dynamic light scattering (DLS) to confirm a mean particle size of approximately 50 nm, and fibril fragmentation was verified by TEM imaging following sonication (Supplementary Figure 14B), then aliquoted into sterile microcentrifuge tubes (20–100 µL), and stored at −80 °C. For quality control, the generated PFFs were routinely characterized by DLS using Zetasizer Nano S (Malvern) and TEM imaging, following previously established procedures^56^.

### Stereotaxic Surgery & Injections

Marmosets were anesthetized with ketamine (15-20 mg/kg) mixed with medetomidine (0.025 mg/kg). Stereotaxic coordinates targeting the caudate nucleus and putamen were derived from the Paxinos marmoset brain atlas^18^. All animals received unilateral intracerebral injections into the left hemisphere, with the exception of one animal (Marmoset Ju), which was injected in the right hemisphere. Animals were randomly assigned to receive either PFF or PBS.

At Western University, injections were performed using an integrated stereotactic injector system (Stoelting, automated stereotaxic arm). Burr holes (1.4 mm) were drilled at the target coordinates on the exposed skull. A 50 µL Hamilton syringe (28-gauge needle) was used for infusion. PFFs injections consisted of 25 µL aliquots of 5 mg/mL marmoset αSyn fibrils, while control injections consisted of 25 µL aliquots of 0.1 M PBS. All injections were delivered at a rate of 1 µL/min and targeted to the caudate nucleus (anterior +11.0 mm, lateral 3.0 mm, depth 6.0 mm) and putamen (anterior +11.0 mm, lateral 5.5 mm, depth 6.5 mm). For each site, the needle was lowered to the cortical surface to define a reference position and then advanced to target depth. Tissue was allowed to settle for 5 minutes prior to infusion onset. After completion of each injection, a 5-minute waiting period was observed before slow needle retraction, and the second site was subsequently targeted following the same procedure. Burr holes were sealed with bone wax upon completion.

In addition, all marmosets undergoing MRI and fMRI procedures at Western University were implanted, in the same surgical session, with a PEEK head post to enable head fixation during imaging. Head post implantation followed previously established procedures^57^. Briefly, following completion of the injection procedure and sealing of burr holes with bone wax, the exposed skull was prepared with adhesive resin (All-Bond Universal, Bisco), and the head post was secured using a resin composite (Core-Flo DC Lite, Bisco). Physiological parameters were continuously monitored throughout surgery. Animals recovered for two weeks post-operatively and were subsequently acclimated to the head-fixation system over approximately three weeks in a mock MRI environment.

At McGill University, injections followed a robot-assisted stereotaxic procedure (Brainsight system, Rogue Research) integrated with a stereotactic frame. A scalp incision was made at the vertex, and the skull was exposed, cleaned of soft tissue, and prepared for optical registration. A laser-based 3D registration grid was used to align the Brainsight system prior to surgery. Burr holes were drilled using a robotic drill arm under veterinary supervision, and completeness of skull penetration was verified manually. Injections were performed using a 100 µL Hamilton syringe fitted with a 22-gauge needle, mounted on the robotic injector arm. The injector was lowered under real-time robotic guidance to the target coordinates, aligned with the drilled aperture, and injections were performed following the same infusion parameters as at Western University (1 µL/min; 25 µL per site), targeting the same striatal coordinates.

### Perfusion and Tissue Fixation

Animals were perfused at different timepoints following stereotaxic injection of αSyn PFF or PBS in order to characterize the temporal progression and spreading of pathology, as well as to enable direct comparison with control animals. Detailed perfusion timepoints for each subject are provided in Supplementary Table 2. Animals were anesthetized via an intramuscular injection of ketamine (15–20 mg/kg) combined with medetomidine (0.025 mg/kg). Following sedation, anesthesia was maintained using isoflurane (1–2%) delivered with oxygen at a flow rate of 1.5–2 L/min. Isoflurane concentration was gradually increased until a surgical plane of anesthesia was achieved. An intravenous catheter was inserted, and a loading dose of propofol (2–5 mg/kg) was administered immediately prior to perfusion. Physiological parameters, including heart rate and arterial oxygen saturation (SPO₂), were continuously monitored using a pulse oximeter, with additional monitoring via stethoscope and visual assessment of respiration rate. An absence of reflexes was confirmed using a toe pinch before initiating perfusion. The thoracic cavity was opened to expose the heart, and a perfusion needle connected to the perfusion pump was inserted into the lower left ventricle. The right atrium was incised to allow efflux of perfusate. For fixed brain perfusion, heparinized saline (0.3 mL heparin per 1 L saline) was first run through (300–400 mL) until the effluent coming from heart was clear. This was followed by perfusion with 4% paraformaldehyde (PFA) for 5–10 min to fix the tissue. Adequate fixation was confirmed by palpating for stiffness in the neck muscles. Involuntary muscle contractions of the limbs and tail were observed during the initial phase of fixative perfusion, indicating tissue penetration. All fixed brain perfusions were performed either on a downdraft table in a designated post-mortem facility or, if unavailable, under specialized respirators with appropriate chemical cartridges. Following PFA/formalin use, the perfusion room was sealed for 24 h to allow dissipation of residual fumes.

### Immunohistochemistry for detection of a-synuclein pathologies

Brains were post fixed in 4% PFA for 1 week, followed by storage in 1x PBS + 0.02% sodium azide. To preserve cellular structure of brains, brains were cryoprotected in 30% sucrose in 1x PBS + 0.02% sodium azide for 10 days. After fixation, brains were frozen with Optimal Cutting Temperature solution (OCT) (Thermofisher, Cat. 6769006). Whole Brains were mounted on a chilled -20_°_C platform and sectioned coronally using a cryostat (Leica) at 30µm thick sections. Sections were collected into 1x PBS + 0.02% sodium azide and stored at 4_°_C. Free floating sections were mounted onto SuperFrost plus slides and treated to 10mM sodium citrate buffer with 0.05% Tween-20 (pH 6.0) at 94_°_C for 20 mins, followed by cooling to room temperature. Sections were washed in 1 × 10min in 1x tris-buffered saline (TBS), followed by 3 × 10 min washes of tris-buffered saline with tween-20 (TBS-T) containing 0.2% Triton buffer, and blocked for 1 hr in diluted normal serum blocking buffer (5% donkey serum, 2% goat serum) in 0.3% TBS-T at room temperature. Primary antibodies, 1:600 rabbit anti-phosphorylated αSyn at serine 129 (Abcam, ab51253, Lot# 1108974-55) and 1:500 mouse anti-tyrosine hydroxylase (Biolegend, 818001, Lot# B421098) was incubated in blocking buffer overnight at 4_°_C. Sections were washed 3 × 10 min in TBS-T, followed a by the incubation secondary antibodies, 1:1000 Alexa647 conjugated donkey anti-rabbit IgG and 1:1000 Alexa555 conjugated donkey anti-mouse IgG for 2 hours in blocking buffer at room temperature. Further washing using TBS-T was done, followed by a 10 min incubation in 1:1000 Hoechst in TBS. Final washes in TBS were done and sections were cover slipped with immu-mount (Epredida, Cat. 9990402) mounting medium. Images were taken with a Leica MICA microscope, and Leica Stellaris5 Confocal microscope. Using the Leica MICA system, images were acquired with the 405 nm and 638 nm excitation channels to visualize Hoechst and αSyn, respectively. 20x magnification tile scans were collected to capture regional staining patterns across each section. For dual-label pS129 and TH imaging in the substantia nigra (Supplementary Figure 8), four adjacent 40x tile scans were acquired and merged to generate a composite image covering the medial portion of the substantia nigra at +5.0 mm from the interaural line (as defined in the Paxinos marmoset brain atlas^18^), at each post-injection timepoint examined (2, 5, 7, 11, and 13 months). Confocal z-stacks acquired at 40x magnification had a total stack depth of 24.11µm, generating 81 optical slices. Higher-resolution imaging was performed on the Leica Stellaris5 confocal microscope, using the same excitation wavelengths (405 nm for Hoechst, and 638 nm for αSyn). Confocal z-stacks were acquired at 63x magnification with a total stack depth of 27.23µm, generating 92 optical slices, which were processed into maximum-intensity projections for visualization and analysis. For Figure 2, 63x images were acquired from the caudate and putamen at the injection site (+11.0 mm from the interaural line) and from the substantia nigra and hippocampus (CA1) at +4.5 mm from the interaural line.

### Structural MRI and Resting-State fMRI Acquisition

A total of 11 marmosets underwent structural MRI scanning, including 6 animals in the PFF group (2 females) and 5 in the PBS group (1 female). Of these, three animals (2 PFF, 1 PBS) were scanned at McGill University, while the remaining animals were scanned at Western University. Detailed information on scanning availability for each subject is provided in Supplementary Table 2.

At Western University, awake animals were positioned in a custom 3D-printed sphinx-style chair with head fixation achieved via an implanted PEEK head post^57,58^. A black fixation dot on a grey background was rear-projected onto a screen located 119 cm from the eyes (Sony VPL-FE40 LCD projector and front-surface mirror) to minimize eye movements and reduce nystagmus, consistent with our previous marmoset fMRI studies. Silicone earplugs were inserted into the ear canals for acoustic protection, and an MRI-compatible camera (Model 12M-i, 60 Hz, MRC Systems GmbH) was used for continuous monitoring of the animal’s state during scanning.

At McGill University, animals were similarly seated in a custom 3D-printed sphinx-style chair; however, head stabilization was achieved using individualized 3D-printed helmets based on CT-derived head models rather than implanted head posts. The same visual fixation stimulus was presented to minimize eye movements. Animal state was monitored using an MRI-compatible camera (Model 12M-i, 60 Hz, MRC Systems GmbH).

At both sites, T2-weighted anatomical images were acquired longitudinally at multiple timepoints: baseline, and 3, 6, 8, and 11 months post-injection. Baseline scans differed slightly between sites: at McGill University, baseline images were acquired prior to injection, whereas at Western University, baseline scans were acquired approximately 1-month post-injection to allow for recovery from head post implantation and completion of behavioral training.

At Western University, imaging was performed on a 9.4T system (31-cm bore Varian magnet interfaced with a Bruker Avance NEO console) using a custom 15-cm gradient coil and an eight-channel receive coil within a quadrature birdcage transmit coil^58^. T2-weighted images were acquired with the following parameters: TR = 7 s, TE = 39 ms, flip angle = 160°, slice thickness = 0.5 mm, in-plane resolution = 0.133 × 0.133 mm², and GRAPPA factor = 2. At McGill University, imaging was performed on a 7T whole-body MRI system (MAGNETOM Terra, Siemens Healthineers) using a 24-rung quadrature birdcage transmit coil (15 cm diameter) and a custom 8-channel phased-array receive coil integrated into the head stabilization helmet^59^. T2-weighted images were acquired with the following parameters: TR = 3 s, TE = 515 ms, field of view = 70 × 59.1 mm², isotropic voxel size = 0.2 mm, bandwidth = 0.35 kHz/pixel, and GRAPPA factor = 2.

Resting-state fMRI (rs-fMRI) data were acquired exclusively at Western University using the same scanner and coil configuration described above. Eight animals were included in the fMRI cohort (4 PFF, including 1 female; 4 PBS, including 1 female). Functional images were acquired using a gradient-echo, single-shot echo planar imaging (EPI) sequence with the following parameters: TR = 1.5 s, TE = 15 ms, flip angle = 40°, field of view = 64 × 48 mm², isotropic voxel size = 0.5 mm, bandwidth = 400 kHz, and GRAPPA factor = 2.

Each fMRI session consisted of four 15-minute resting-state runs. Animals were scanned at baseline (approximately 1-month post-injection), as well as during early (3–6 months) and late (8–11 months) post-injection windows. A total of two sessions were acquired at baseline and four sessions during each post-injection window. Due to early endpoint for histological analysis, one PFF-injected female (marmoset Ju) contributed data only at baseline and during the early time window. Detailed information on scanning availability for each subject is provided in Supplementary Table 2.

### Structural MRI preprocessing and deformation-based morphometry

All anatomical images were preprocessed using a standardized pipeline implemented in a combination of FSL^60^, ANTs^61^, and AFNI^62^ tools. Preprocessing steps were identical for data acquired at both sites.

First, images were reoriented to a common reference space to ensure compatibility with subsequent processing steps. Intensity inhomogeneity was then corrected using N4 bias field correction. Each image was subsequently registered to a study-specific template based on the Marmoset Brain Mapping (MBM) atlas^19^ using affine transformations. A brain mask derived from the template was transformed into individual subject space and applied to remove non-brain tissue. Finally, images and corresponding masks were resampled to the template grid to ensure consistent spatial resolution and alignment across subjects and timepoints.

Following preprocessing, deformation-based morphometry (DBM) analyses were performed using a two-level registration framework adapted from established pipelines^63^. Briefly, subject-specific templates were first generated from longitudinal data, and these were then used to construct a group-level template (including both PFF and PBS samples) representing the study population. All individual images were nonlinearly registered to the group template using symmetric normalization. Voxelwise estimates of local volumetric change were derived from the deformation fields by computing the logarithm of the Jacobian determinant at each voxel. These log-Jacobian maps reflect regional expansions and contractions relative to the group template. To improve signal-to-noise ratio and account for residual anatomical variability, log-Jacobian maps were spatially smoothed using a Gaussian kernel (σ = 1 voxel) and registered to the MBM template^19^ prior to statistical analysis.

Statistical analyses of voxelwise log-Jacobian maps were performed using a linear mixed-effects (LME) model implemented in AFNI (3dLMEr). The model included fixed effects of Group (PFF vs PBS), Timepoint (treated as a continuous variable), and their interaction, with Monkey included as a random intercept to account for repeated measures. This approach enabled the assessment of both group differences and longitudinal trajectories of structural change.

To interrogate specific effects, general linear tests were defined to evaluate: (i) the main effect of Group (PFF vs PBS), (ii) the within-group longitudinal slope for each condition (PFF and PBS separately), and (iii) the difference in slopes between groups (Group*Timepoint interaction), reflecting differential progression of structural changes over time. Statistical maps were thresholded using a voxelwise significance level of p<0.001, and cluster-level correction (α<0.05) was applied using a minimum cluster size of 50 contiguous voxels (nearest-neighbor connectivity = 1). This procedure identified clusters showing significant group differences in longitudinal trajectories.

To further characterize these effects, mean log-Jacobian values were extracted from significant clusters for each subject and timepoint. These values were then used to visualize and quantify longitudinal changes, enabling direct comparison of structural trajectories between PFF- and PBS-injected animals.

### Resting-state fMRI preprocessing

Rs-fMRI data were preprocessed using a standardized pipeline combining tools from AFNI^62^, FSL^60^, and ANTs^61^. All preprocessing steps were applied at the single-run level and were identical across subjects.

Functional images were first reoriented to a common reference space to ensure consistency across runs and compatibility with subsequent processing steps. Susceptibility-induced distortions were then corrected using a fieldmap-free approach based on pairs of images acquired with opposite phase-encoding directions (topup, FSL), and the resulting deformation fields were applied to the full time series. Data quality was assessed by computing the fraction of outlier voxels at each timepoint. Transient signal spikes were attenuated using despiking, and slice timing differences were corrected. Head motion was then corrected by rigid-body realignment of all volumes to a reference volume within each run, and motion parameters were estimated for subsequent censoring. A mean functional image was computed for each run following motion correction. Each run was aligned to the corresponding subject-specific T2-weighted anatomical image using affine registration. Tissue-specific masks (white matter and cerebrospinal fluid), derived from anatomical images, were transformed into functional space and used to extract nuisance time series.

Nuisance regression was performed to remove confounding signals, including white matter and cerebrospinal fluid signals. Linear and higher-order polynomial trends were also removed to account for slow signal drifts. Volumes with excessive motion were censored based on a framewise displacement threshold (0.25 mm), including the preceding timepoint. Following regression, data were spatially smoothed using a Gaussian kernel of 1 voxel and temporally filtered using a bandpass filter (0.01–0.1 Hz) to retain low-frequency fluctuations associated with resting-state activity. Preprocessed functional images were skull-stripped with a brain mask derived from the individual anatomical images and then normalized to the MBM^19^ template using transformations derived from anatomical registration.

### Functional Connectivity and seed-based analysis

Functional connectivity matrices were computed using region of interest (ROI)-based correlations of fMRI time series. ROIs were defined based on the Paxinos^18^ parcellation of the MBM atlas^19^, complemented by selected subcortical regions from a dedicated marmoset subcortical atlas^20^ (including, among others, caudate, putamen, thalamus, hippocampal formation, and substantia nigra). For each run, the mean BOLD time course was extracted from all ROIs. Time courses were averaged across runs within each session and subsequently across sessions within each timepoint, yielding a single representative time series per ROI for each subject and timepoint. To ensure anatomical consistency across subjects, the single animal injected in the right hemisphere (marmoset Ju) was flipped along the left–right axis after preprocessing. As a result, all analyses were performed in a common reference frame where the injection site was aligned across animals. Accordingly, connectivity results are described in terms of *ipsilateral* and *contralateral* relative to the injection site, rather than anatomical left and right.

Pairwise functional connectivity between ROIs was quantified using Pearson correlation coefficients, which were subsequently Fisher z-transformed for statistical analyses. Individual correlation matrices were computed for within-hemisphere and interhemispheric connections and combined into a single matrix while preserving anatomical ordering. At the group level, connectivity matrices were obtained by averaging subject-level matrices element-wise.

Baseline functional connectivity matrices were analyzed separately for the two experimental groups (PBS and PFF), while maintaining a distinction between ipsilateral, contralateral, and inter-hemispheric connectivity. For each subject, global mean functional connectivity was computed as the average of all pairwise correlation values (excluding the diagonal for intra-hemispheric matrices). This metric captures both the overall strength of connectivity and its sign (positive vs negative). In addition, similarity between connectivity patterns was quantified using the Frobenius distance. Pairwise Frobenius distances were computed between all pairs of subjects as the square root of the sum of squared differences between corresponding elements of their connectivity matrices. Distances were classified as within-group or between-group comparisons. These analyses were performed both on the full connectivity matrices and on a subset restricted to regions belonging to the frontal cluster, which was selected as the cluster showing the largest decline in connectivity following PFF injection in the longitudinal analyses. Group differences in global functional connectivity (PBS vs PFF) and in Frobenius distances (within-group vs between-group) were assessed using unpaired t-tests, to evaluate potential baseline differences in functional connectivity independent of the injection.

To investigate longitudinal changes in functional connectivity, delta (Δ) connectivity matrices were then computed for each subject by subtracting baseline connectivity from subsequent time windows (early − baseline; late − baseline), using the Fisher z-transformed matrices. From these Δ matrices, two complementary metrics were derived. First, the mean signed change (MSC) was computed as the average of all matrix elements, providing an estimate of the overall direction of connectivity changes (increase vs decrease). Second, Frobenius distances were computed to quantify the magnitude of changes in connectivity patterns. These metrics were calculated, as previously described, for full-brain matrices (separately for ipsilateral, contralateral, and inter-hemispheric connectivity) and for the frontal cluster. Within the frontal cluster, connectivity was further subdivided into frontal-to-frontal (within-cluster) and frontal-to-rest (between-cluster) connections. Statistical evaluation of longitudinal effects was performed using LME models with Sample (PBS vs PFF) and Timepoint as fixed effects and Monkey as a random intercept.

In addition to whole-brain connectivity, seed-based analyses were performed using selected ROIs, including the caudate nucleus (injection site), the hippocampal formation (showing high αSyn burden in histology), a frontal cluster encompassing areas 46 and 8 (identified from whole-brain connectivity reductions) and the mediodorsal (MD) thalamus. The inclusion of this last one was motivated by its central role in thalamo-cortical circuits. The MD thalamus is strongly interconnected with prefrontal regions, particularly areas 8 and 46, and is critically involved in cognitive and executive functions^35,36,64,65^. As such, it represents a key hub for assessing whether subcortical-cortical communication within fronto-striato-thalamic circuits is affected.

For each seed region, voxelwise functional connectivity maps were computed by correlating the seed time course with all brain voxels for each run. Resulting maps were Fisher z-transformed and subsequently averaged within subject in two steps: first across runs belonging to the same session, and then across sessions belonging to the same timepoint (baseline, early window, late window), yielding one connectivity map per subject and timepoint. Group-level statistical analysis was performed using a voxelwise LME model implemented in AFNI (3dLMEr). The model included Sample (PBS vs PFF) and Timepoint as fixed effects and Monkey as a random intercept. Timepoint was modelled as a quantitative variable to estimate longitudinal trends. From this model, we derived contrasts assessing (i) the main effect of group, (ii) within-group slopes across time (PBS and PFF separately), and (iii) differences in slopes between groups. Maps of slope differences (PFF vs PBS across timepoints) were thresholded using a voxelwise significance level of p<0.001, and cluster-level correction (α<0.05) was applied using a minimum cluster size of 50 contiguous voxels (nearest-neighbor connectivity = 1); this allowed to visualize regions showing differential longitudinal changes in connectivity with the seed. This approach provides a more spatially explicit representation of connectivity changes compared to matrix-based analyses, particularly when focusing on the network profile of individual regions.

### Regional homogeneity (ReHo) analysis

For each resting-state fMRI run, voxelwise maps of ReHo were computed to quantify the local synchronization of the BOLD signal. ReHo measures the similarity of the time series of a given voxel with those of its neighbouring voxels, typically using Kendall’s coefficient of concordance, and is considered an index of local functional coherence^21^. Given the sensitivity of ReHo to noise and local signal characteristics, additional precautions were taken during preprocessing. After nuisance regression and motion censoring, and prior to spatial smoothing, bandpass filtering, and normalization to template space, each run was masked to exclude white matter and cerebrospinal fluid signals. ReHo maps were then computed in native space for each run. Subsequently, these maps were spatially normalized to the template and smoothed using a 2 mm full-width at half-maximum Gaussian kernel.

Run-level ReHo maps were first averaged within session and then across sessions belonging to the same timepoint (baseline, early window, late window), while keeping individual animals separate. This resulted in one ReHo map per animal per timepoint. Voxelwise statistical analysis was performed using a linear mixed-effects LME model implemented in AFNI (3dLMEr), with Group (PBS vs PFF) and Timepoint as fixed effects and subject as a random intercept. Specific contrasts were defined to assess group differences at each timepoint, within-group longitudinal changes, and Group*Timepoint interactions. Statistical maps corresponding to the interaction effects were cluster-corrected using a voxelwise significance level of p<0.001, and a minimum cluster size of 50 contiguous voxels (nearest-neighbor connectivity = 1, α<0.05).

To further characterize significant interaction effects, clusters identified at the late timepoint (11 months; PFF vs PBS interaction relative to baseline) were used as ROIs. Within these clusters, ReHo values were extracted for each monkey, session, and timepoint. Values were then averaged within timepoint and normalized relative to baseline. These data were used to visualize both observed trajectories and model-predicted effects, enabling a detailed characterization of the temporal dynamics underlying the interaction effects.

### Actimetry

Motor activity and sleep patterns were monitored using the CamNtech Nano veterinary actigraph (CamNtech Ltd., UK) at both sites. At McGill, the device was attached to a collar worn by the animal, whereas at Western it was mounted on a custom holder fixed to the headpost. This latter configuration allowed for increased sensitivity to head movements, even in the absence of overt body motion. Actimetry recordings were collected approximately every two months, starting one month after injection. A total of 11 marmosets were included (Western: *n*=8, 4 PFF and 4 PBS; McGill: *n*=3, 2 PFF), with sex distribution detailed in Supplementary Table 2. To minimize variability related to experimental procedures, recordings were performed over weekends only. Devices were placed on Friday evening and initiated one hour prior to lights-off (18:00 h at Western, 19:00 h at McGill), and removed on Monday morning, one hour after lights-on (8:00 h at Western, 9:00 h at McGill). During these periods, animals were maintained under standard housing conditions with only routine feeding, avoiding potential confounds from scanning or behavioural testing. Activity data were acquired at a temporal resolution of 5-second epochs.

Raw actigraphy data were preprocessed to classify each time point as sleep or wake based on an activity threshold (epochs with <30 motor acts were considered as sleep), followed by identification of sustained bouts to reduce spurious transitions. From these processed time series, a set of behavioural metrics was derived separately for night and day periods.

Night-time metrics included: (i) **Total sleep duration**, defined as the total time spent in the sleep state during the night; (ii) **Sleep latency**, defined as the time from lights-off to the first sustained sleep bout; and (iii) **Transitions**, defined as the number of state changes between sleep and wake across the night, providing an index of sleep fragmentation. Daytime metrics included: (iv) **Mean activity**, defined as the average activity level across the day; (v) **Number of naps**, corresponding to the number of discrete sleep bouts occurring during the day; and (vi) **Nap duration**, defined as the cumulative time spent asleep during the day.

These metrics were computed for each recording session and subsequently averaged within timepoints for each animal, enabling longitudinal assessment of motor activity and sleep alterations across groups. To assess longitudinal changes in actimetry-derived metrics, statistical analyses were performed using LME models, where Group (PBS vs PFF) and Timepoint were treated as fixed effects, and Monkey was included as a random intercept to account for repeated measures within subjects. For each model, main effects of Group and Timepoint, as well as their interaction, were evaluated using analysis of variance (ANOVA). When a significant main effect of Timepoint was observed, post-hoc pairwise comparisons between timepoints were performed using estimated marginal means with Tukey correction for multiple comparisons. In the presence of a significant Group*Timepoint interaction, post-hoc comparisons were conducted to assess group differences at each timepoint, again using Tukey-adjusted contrasts. Estimated marginal means were also computed to characterize group-level trajectories over time and to support visualization of the longitudinal profiles of each metric.

### Pairwise Visual Discrimination (PVD) Task

Cognitive performance was assessed using a touchscreen-based PVD task. A total of 10 marmosets (PBS: *n*=5, 1 female; PFF: *n*=5, 1 female; see Supplementary Table 2) were tested longitudinally at baseline (pre-injection) and at 5, 7, 9, and 11 months post-injection. Due to planned experimental endpoints for histological analyses, the number of animals decreased over time, and the final timepoint (11 months) should be considered exploratory (2 PFF and 1 PBS).

All sessions were conducted in the home cage using a custom-built touchscreen apparatus directly attachable to the cage. The setup (see Supplementary Figure 15A) consisted of a plexiglass testing box (27.5 × 25 × 20 cm), with an opaque floor and transparent walls. One side featured a metal docking plate and sliding door to allow voluntary access from the home cage. On the opposite side, an infrared 10.4-inch touchscreen (Lafayette Instrument) was mounted and connected to a Raspberry Pi 3 running a custom Python-based control software^66^. A reward spout, positioned centrally in front of the screen at a height of 13 cm and made of brass tubing, delivered liquid reward via an infusion pump (model NE-510, New Era Pump Systems).

The PVD task was adapted from established touchscreen paradigms developed by the Bussey–Saksida group^67–70^ and previously adopted also for marmosets^37^. In each trial, two black and white visual stimuli (squared, 5 cm) were presented simultaneously on the left and right side of the screen, separated by 4 cm and positioned around the reward spout (see Supplementary Figure 15B for an example of the visual stimuli used in the task; see Supplementary Video 1 for an example of performance). Each session consisted of a maximum of 30 trials and lasted no longer than 30 minutes, although animals typically completed sessions within 15 minutes. Testing was conducted in the morning prior to feeding, without food or water deprivation.

In the first phase of the task (learning phase), one of the stimuli (S+) was associated with the reward, while the second stimulus (S-) was not. In this phase, animals were required to reach a performance criterion of ≥90% correct responses over the last 50 trials (aggregated across sessions). Upon reaching criterion, the reversal phase was initiated by switching the stimulus–reward contingencies (the initially unconditioned S- became the rewarded stimulus, and vice versa for the S+). The task was considered complete once the same performance criterion was achieved again during reversal. A unique pair of stimuli was used at each timepoint to prevent carry-over of stimulus-specific learning across sessions.

From the behavioural data, the following metrics were computed for each monkey at each timepoint, separately for the learning and reversal phases: (i) number of trials required to reach criterion, (ii) error ratio (including both perseverative and regressive errors), and (iii) task engagement, defined as the number of trials completed within the first three minutes of each session. Errors were classified into *perseverative errors*, defined as consecutive incorrect responses (incorrect–incorrect), and *regressive errors*, defined as incorrect responses following a correct response (correct–incorrect), reflecting failures to maintain or update the learned association.

Statistical analyses were performed using LME models. For each metric, separate models were fitted for the learning and reversal phases, where Group (PBS vs PFF) and Timepoint were treated as fixed effects, and Monkey was included as a random intercept to account for repeated measures. This approach allowed assessment of group differences, longitudinal effects, and their interaction across the different stages of task performance. In the presence of a significant effect, post hoc comparisons were conducted to assess group differences at each timepoint, using FDR-adjusted contrasts.

## DATA AVAILABILITY

All data and analysis code supporting the findings of this study, including pipelines for deformation-based morphometry, resting-state fMRI analyses, and touchscreen and actimetry analyses, are publicly available on Zenodo (Repository Draft for Reviewers).

## ACKNOWLEDGMENTS

We thank the animal care staff at both Western University and at the Neuro, McGill University for their support throughout the study. We are particularly grateful to Emily Truscott, Jessica Hutta and Fernando Chaurand for veterinary care of the animals, and to Sarah Brennan and Shehrbano Farooq Khan for their invaluable assistance in facilitating coordination and communication between the two sites. We thank Cheryl Vander Tuin, Whitney Froese, Hannah Pettypiece, David Everling, Miranda Bellyou, Tyler Cook, Margaux Teil, Dominique Bédard, Daniel Hau-Aquino, Valerie Comtois and Cathy Hunt for animal care and preparation; Dr. Alex Li, David Costa and Ron Postuma for assistance with MRI scanning; Dr. Kyle Gilbert, Peter Zeman, Dr. Pedram Yazdanbakhsh and Marcus J. Couch for coil design and development; Dr. Amr Eed and Dr. Gabriel Devenyi for assistance with statistical analyses and pipeline development; Dr. Raymond K. Wong and Dr. Janahan Selvanayagam for assistance with touchscreen training and code development; and Dr. Ywlianne da Silva, Dr. Kelly Summers, Dr. Shahrzad Bahrampour and Dr. Rodrigo Sandoval for assistance with immunohistochemistry procedures.

This research was enabled in part by computational resources and support provided by Compute Ontario and the Digital Research Alliance of Canada through the Trillium and Nibi clusters.

## AUTHORS CONTRIBUTIONS

A.Z., S.E., J.C.C., R.S.M., M.A.M.P., T.J.B., L.M.S., and V.P. conceived and designed the study. W.L., I.S., and T.M.D. produced the marmoset αSyn pre-formed fibrils. K.J., P.H. and A.Z. performed the surgeries. J.S., J.Y., and C.D. performed immunohistochemistry experiments. A.Z., J.Y., A.D., and M.G. conducted structural MRI acquisition and awake imaging training. A.Z. and A.D. acquired resting-state fMRI data. A.Z. developed the MRI and fMRI preprocessing pipelines and performed deformation-based morphometry, functional connectivity, seed-based connectivity, and regional homogeneity analyses. A.Z., J.S., C.D. and J.Y. acquired actimetry data, and A.Z. analyzed the actimetry data. A.Z. acquired and analyzed touchscreen behavioral data and trained the animals. S.J. supervised the project and contributed to study design and data interpretation. A.Z. and J.S. drafted the manuscript. All authors revised and approved the final manuscript.

