## Supplementary Materials for "α-synuclein pathology drives multi-scale reorganization of fronto-limbic networks and cognitive flexibility in the common marmoset"

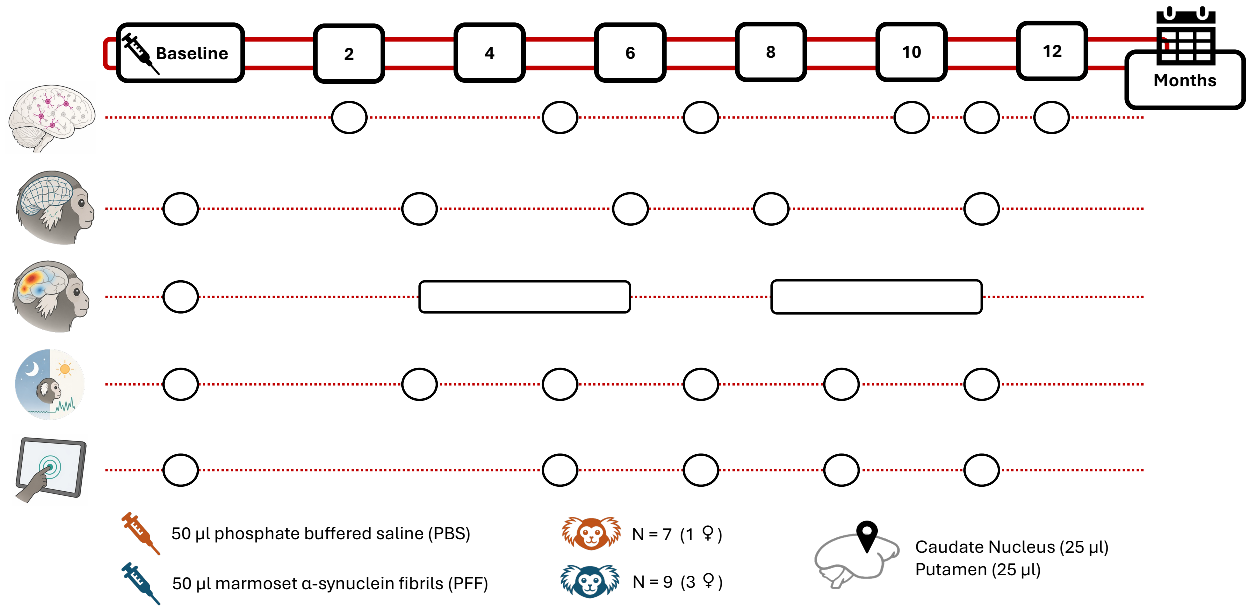


**Supplementary Figure 1. Experimental design and longitudinal data acquisition schedule.** The timeline (top) indicates the post-injection timepoints at which data were acquired (from baseline to 12 months). Rows below the timeline indicate the acquisition schedule for each modality: histology (row 1), structural MRI analyzed by deformation-based morphometry (row 2), resting-state fMRI for functional connectivity matrices, seed-based analyses, and regional homogeneity (row 3), actimetry for continuous sleep and daily motor activity monitoring (row 4), and touchscreen-based pairwise visual discrimination task for cognitive assessment (row 5). Markers positioned along each row indicate the timepoints at which data were collected for the corresponding modality. Bottom panel: animals received unilateral intracerebral injections of either phosphate-buffered saline (PBS; 50 µL, orange) or marmoset-derived αSyn preformed fibrils (PFF; 50 µL, blue). Injections were split across two target structures within the same hemisphere: 25 µL into the caudate nucleus and 25 µL into the putamen. Group sizes are indicated by orange (PBS) and blue (PFF) marmoset symbols.

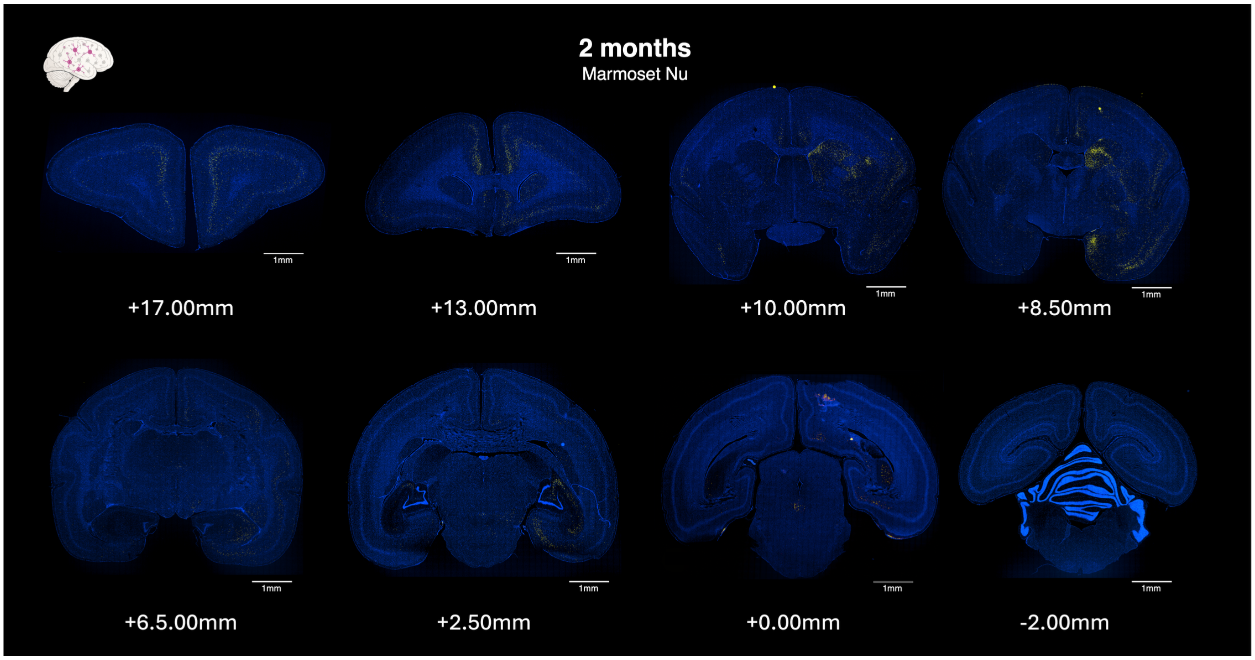

2 months
Marmoset Nu
1mm
+17.00mm
+13.00mm
+10.00mm
+8.50mm
+6.5.00mm
+2.50mm
+0.00mm
-2.00mm


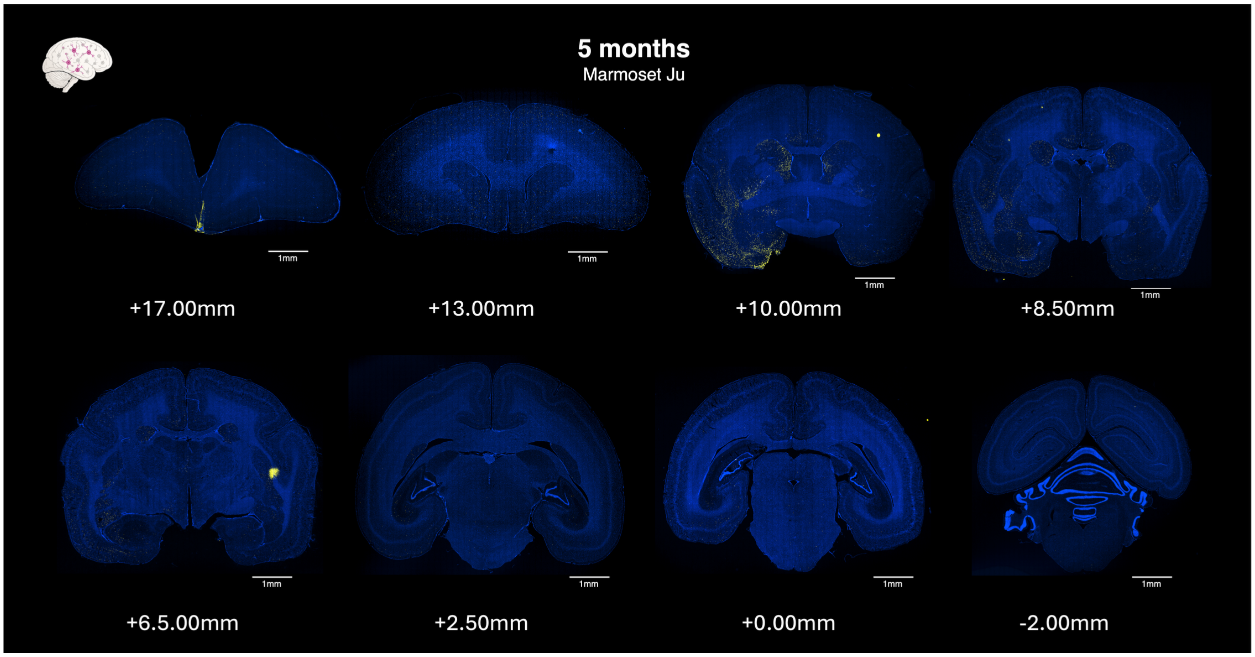

5 months
Marmoset Ju
1mm
+17.00mm
+13.00mm
+10.00mm
+8.50mm
+6.5.00mm
+2.50mm
+0.00mm
-2.00mm

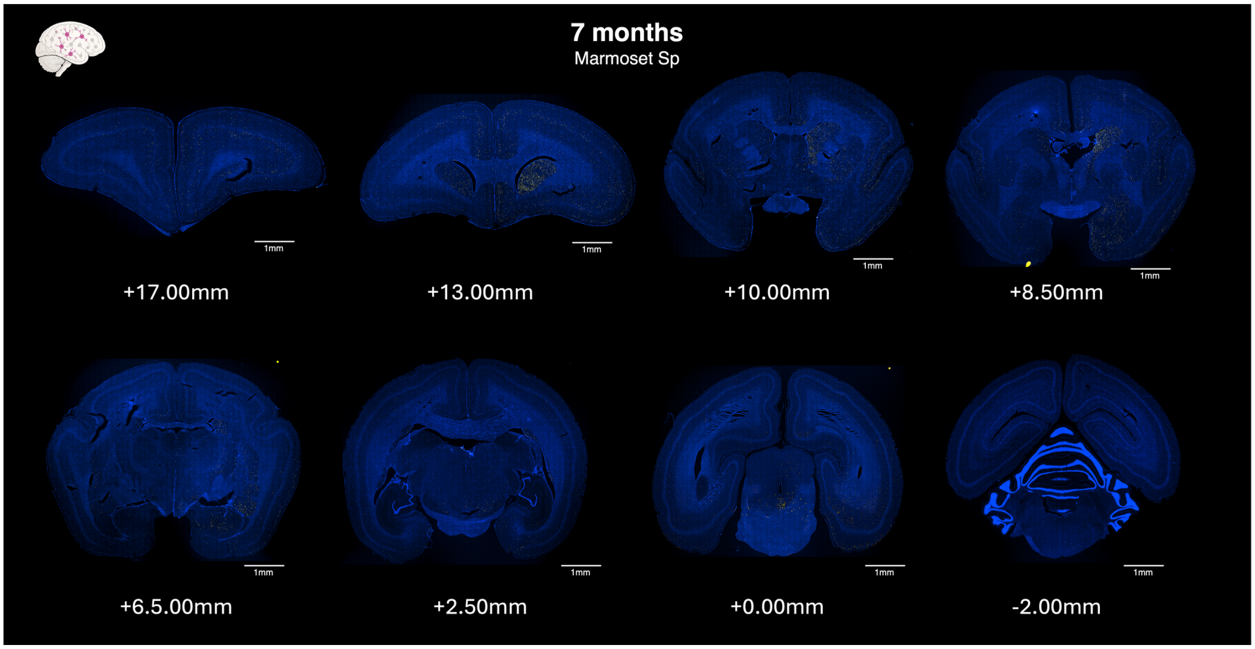
7 months
Marmoset Sp
+17.00mm
+13.00mm
+10.00mm
+8.50mm
+6.5.00mm
+2.50mm
+0.00mm
-2.00mm

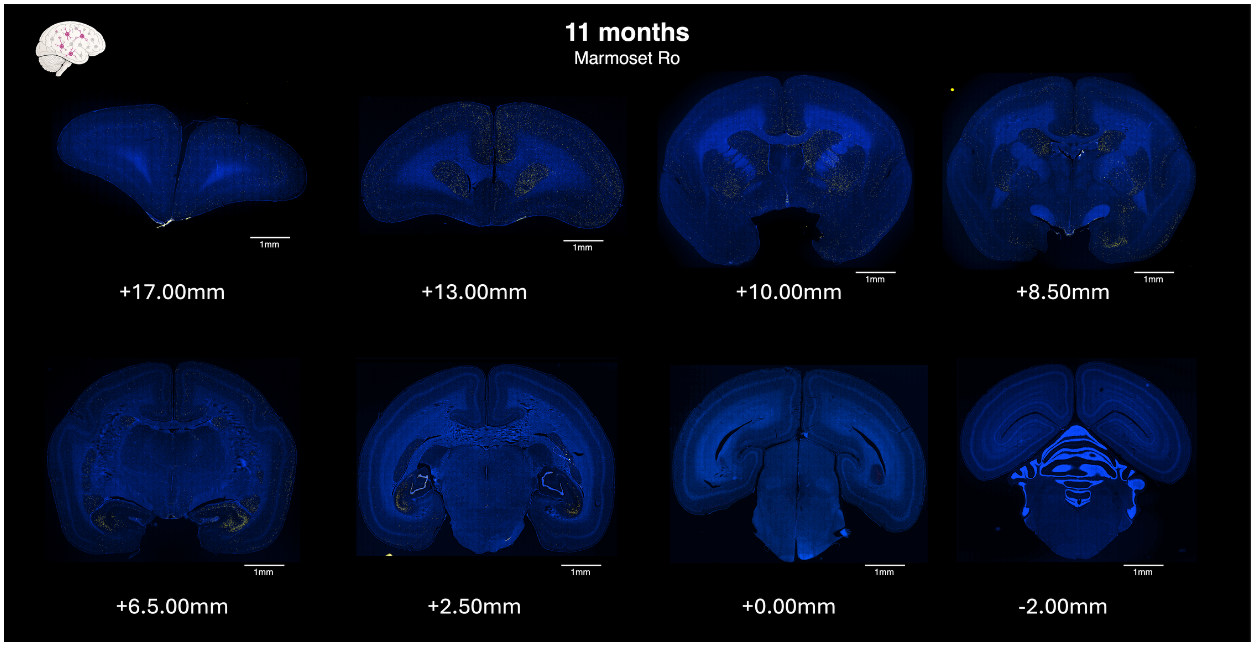
11 months
Marmoset Ro
+17.00mm
+13.00mm
+10.00mm
+8.50mm
+6.5.00mm
+2.50mm
+0.00mm
-2.00mm

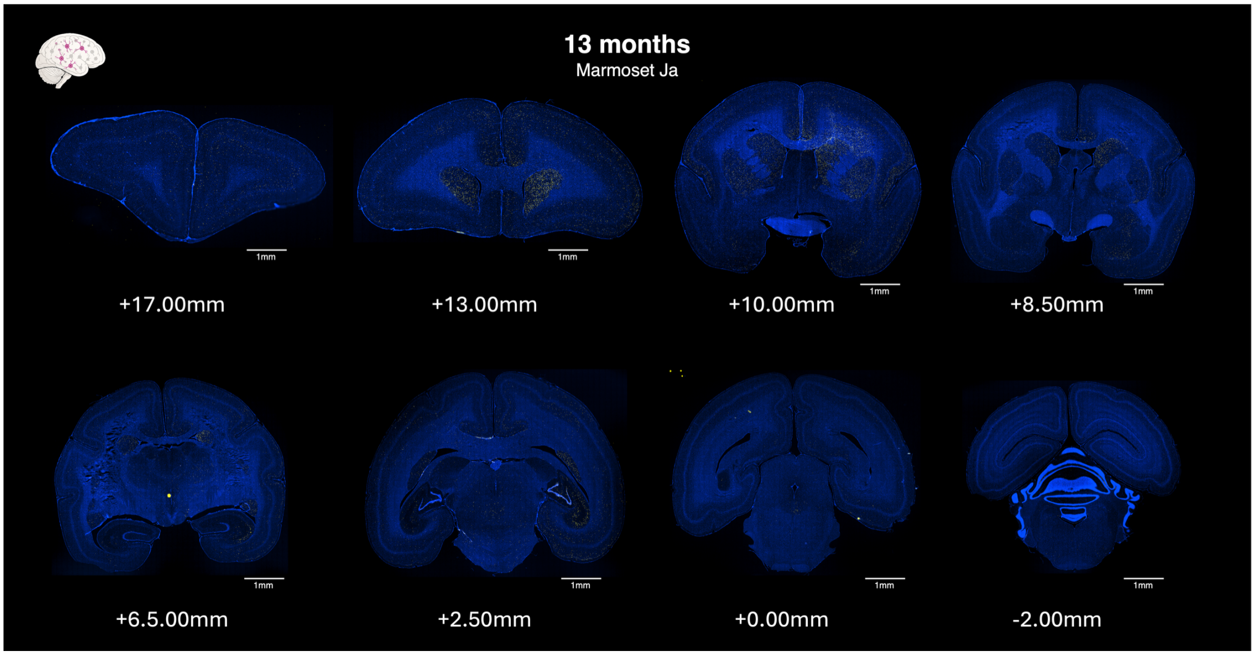


**Supplementary Figures 2–6. Spatiotemporal progression of pS129-positive αSyn pathology across post-injection timepoints.** Coronal sections from PFF-injected marmosets are shown at eight rostrocaudal levels (+17, +13, +10, +8.5, +6.5, +2.5, 0, and −2 mm from the interaural line, as defined in the Paxinos marmoset brain atlas[16]) for five individual animals corresponding to increasing post-injection timepoints: 2 months (Supplementary Figure 2), 5 months (Supplementary Figure 3), 7 months (Supplementary Figure 4), 11 months (Supplementary Figure 5), and 13 months (Supplementary Figure 6). pS129-positive inclusions are shown in magenta; nuclei are counterstained with Hoechst (blue). Together, these figures illustrate the progressive expansion of pathology from early, spatially restricted immunoreactivity to widespread cortical and subcortical involvement across the brain. Scale bars: 1mm.

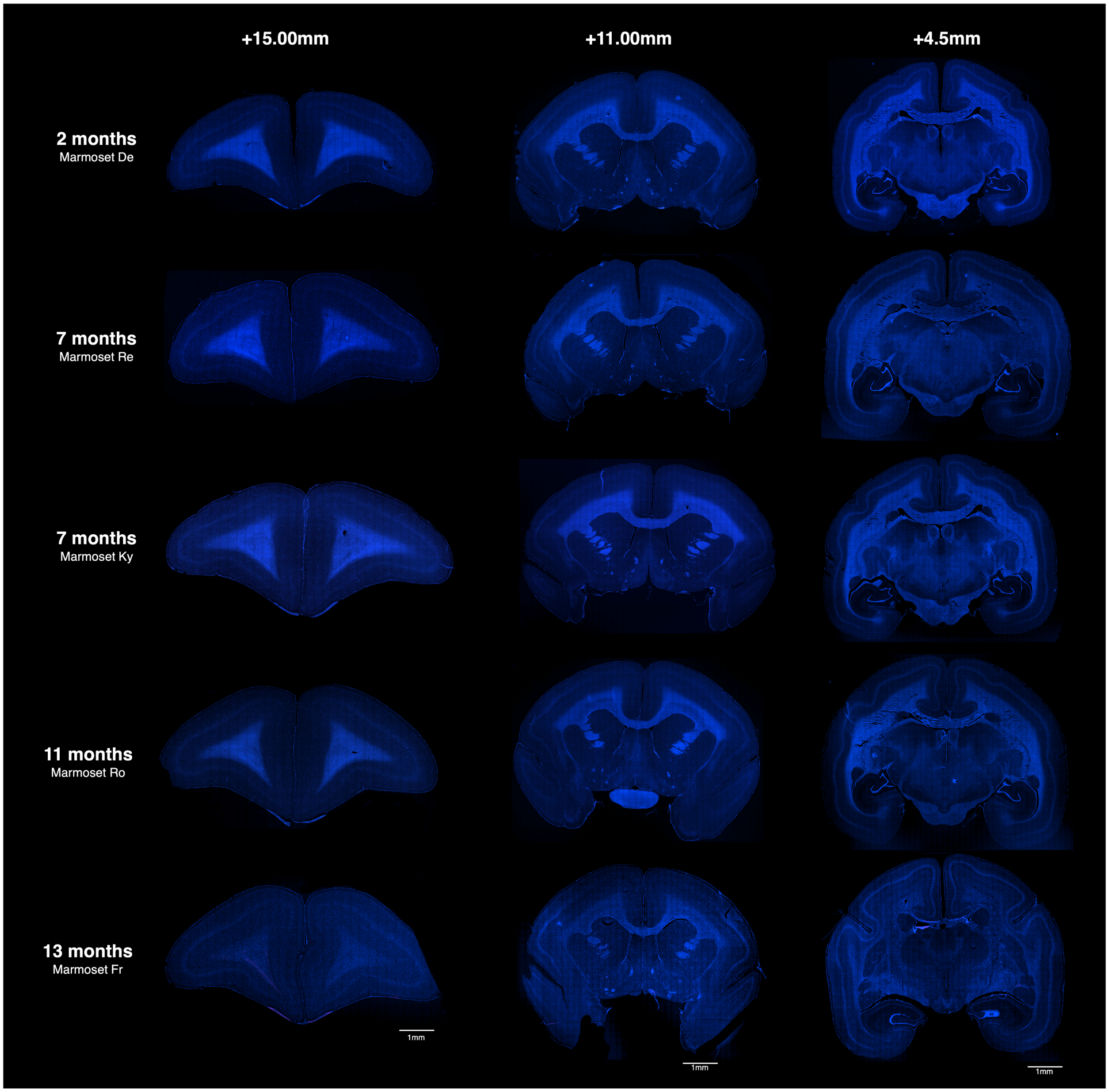


**Supplementary Figure 7. Absence of pS129-positive a-synuclein pathology in PBS-injected control marmosets across post-injection timepoints.** Coronal sections from PBS-injected marmosets are shown at three rostrocaudal levels (+15, +11, and +4.5mm from the interneural line, as defined in the Paxinos marmoset brain atlas[16]) for five individual animals corresponding to increasing post-injection timepoints: (from top to bottom) 2 months, 7, months, 11 months, and 13 months. Sections were immunolabeled for ps129 a-synuclein and counterstained with Hoechst (blue). No detectable ps129-positive inclusions were observed in the PBS-injected animals at any examined timepoint. Scale bars: 1mm.

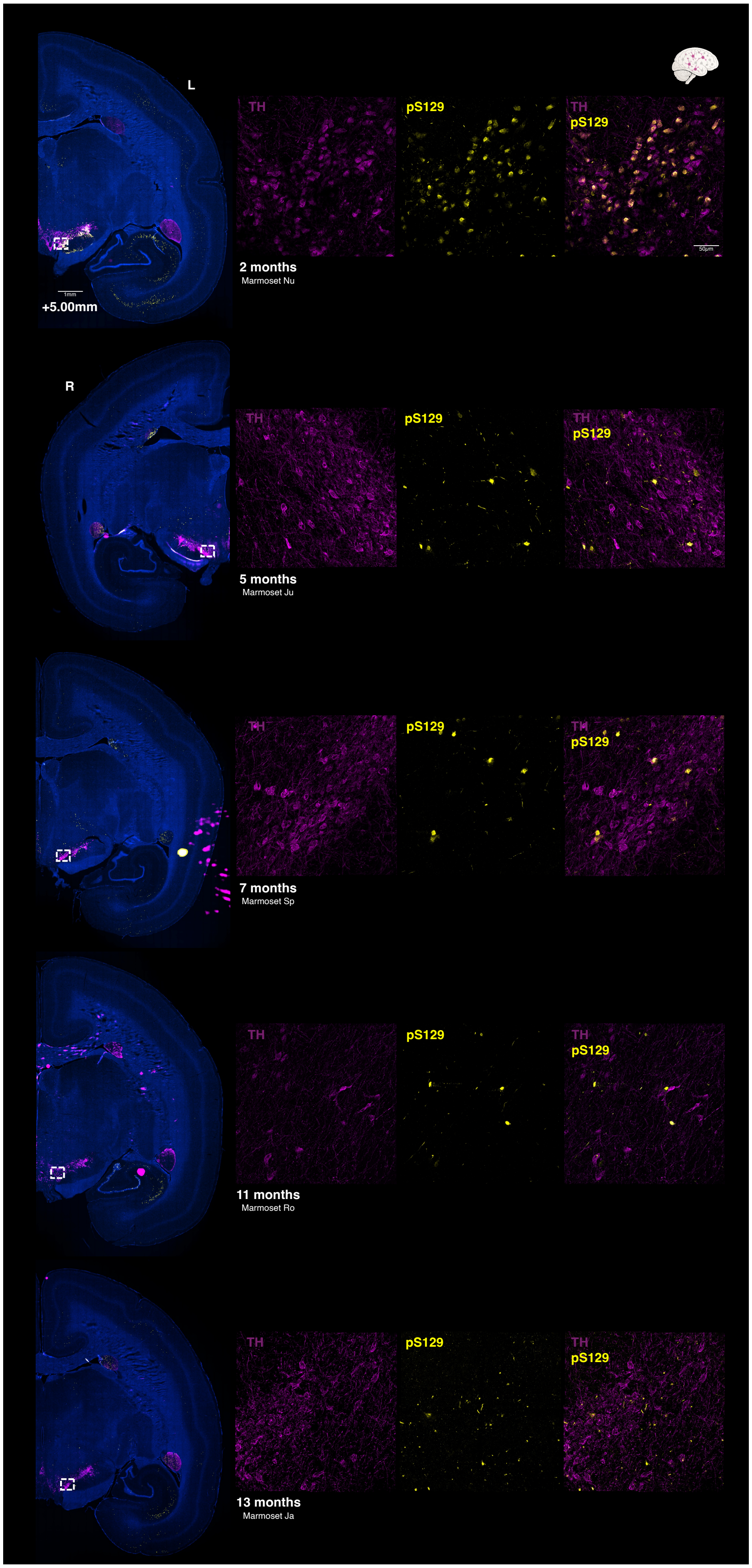
L
TH
pS129
TH
pS129
50μm
2 months
Marmoset Nu
1mm
+5.00mm
R
TH
pS129
TH
pS129
5 months
Marmoset Ju
TH
pS129
TH
pS129
7 months
Marmoset Sp
TH
pS129
TH
pS129
11 months
Marmoset Ro
TH
pS129
TH
pS129
13 months
Marmoset Ja

64

**Supplementary Figure 8. Longitudinal tyrosine hydroxylase (TH) and pS129-αSyn co-immunolabelling in the substantia nigra of PFF-injected marmosets.** Each row corresponds to a single PFF-injected animal at a distinct post-injection timepoint (top to bottom: 2, 5, 7, 11, and 13 months). Left column: 20× tile scan of the injected hemisphere at +5.0 mm from the interaural line, showing the spatial distribution of TH (purple), pS129-αSyn (yellow), and Hoechst nuclear counterstain (blue) across the substantia nigra. The white square indicates the region selected for higher-magnification imaging. Right columns: 40× images of the outlined region, shown separately for TH (left), pS129-αSyn (centre), and the merged channel overlay (right), using the same colour scheme. Together, these images illustrate the progressive evolution of dopaminergic neuron pathology across timepoints, including the sustained preservation of TH immunoreactivity throughout the observation period.

**Functional connectivity – Frobenius Distance and Mean Signed Change.** At baseline, inspection of connectivity matrices revealed regional differences (Supplementary Figure 9), with increased ipsilateral auditory–temporal FC in PFF animals and stronger contralateral frontal FC in PBS animals.

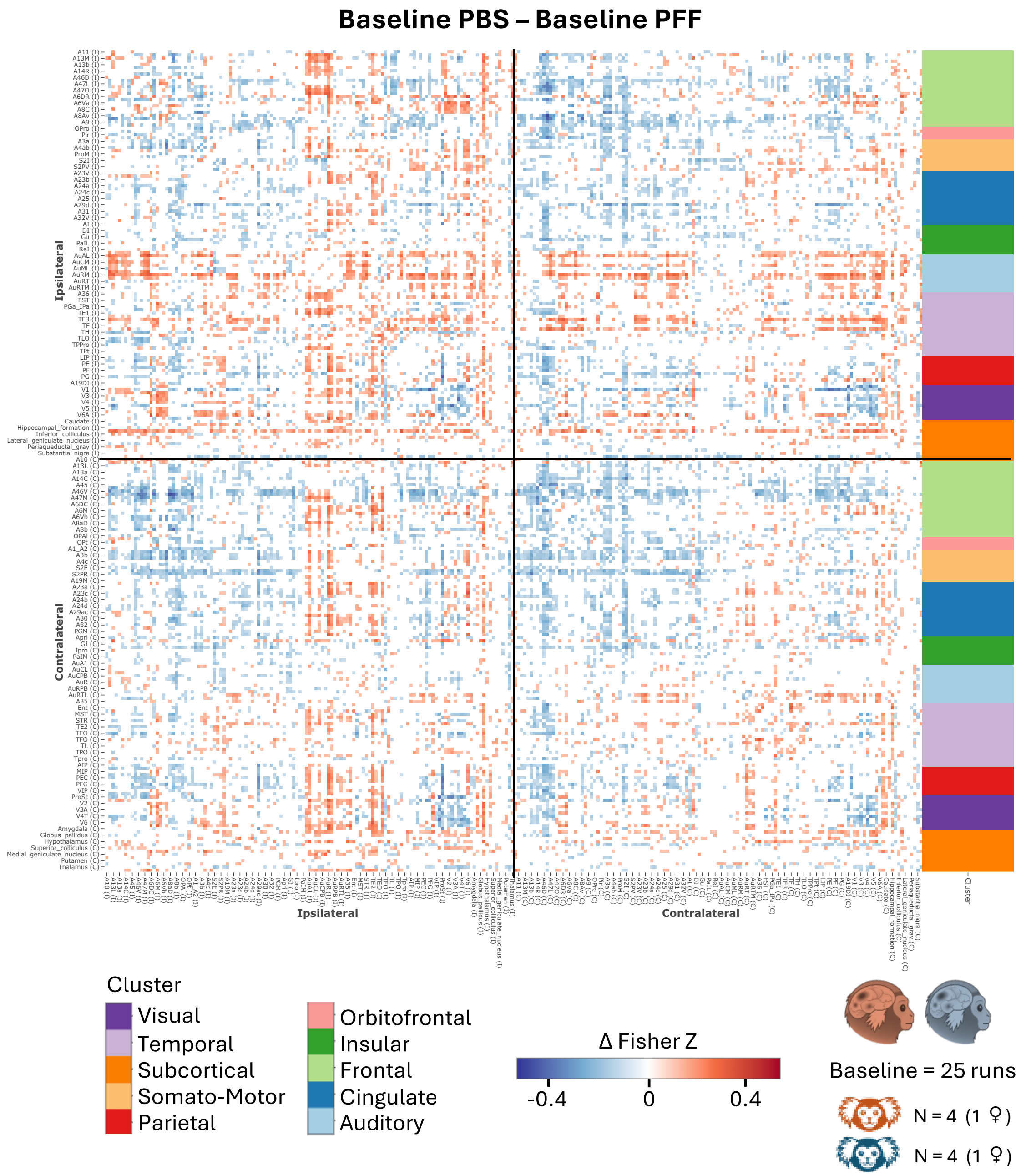


**Supplementary Figure 9. Difference in functional connectivity between groups at baseline.** The matrix shows the direct group difference in functional connectivity at baseline (PFF minus PBS, in Fisher's z units). The matrix is organized into four quadrants reflecting ipsilateral (top left), contralateral (bottom right), and interhemispheric (top right and bottom left, mirrored) connectivity. Blue colors indicate stronger connectivity in PFF animals; red colors indicate stronger connectivity in PBS animals. Colored bars along the matrix y-axis denote anatomical clusters (e.g., light green, frontal regions; pink, orbitofrontal regions) to aid visual interpretation. Regions of interest were drawn from the Paxinos parcellation[18] of the NIH marmoset brain atlas[19], supplemented with subcortical structures from the Subcortical Atlas of the Marmoset[20]. For clarity, odd and even region labels are displayed on the x- and y-axes, respectively.

However, the PFF and PBS groups did not differ in global FC metrics, including global mean FC and Frobenius distance between individuals (all $p>0.05$; Supplementary Table 1), indicating comparable overall network organization prior to injection.

| Measure | Timepoint | Matrix | Connectivity | T | DF | P | Mean PFF | Mean PBS |
|---|---|---|---|---|---|---|---|---|
| GFC | Baseline | Ipsilateral | Whole Matrix | -0.29 | 5.27 | 0.78 | 0.22 | 0.21 |
| GFC | Baseline | Contralateral | Whole Matrix | 0.63 | 5.71 | 0.74 | 0.22 | 0.23 |
| GFC | Baseline | Interhemispheric | Whole Matrix | 0.35 | 6 | 0.55 | 0.20 | 0.21 |
| GFC | Baseline | Ipsilateral | Front to Front | 0.13 | 4.71 | 0.90 | 0.34 | 0.35 |
| GFC | Baseline | Contralateral | Front to Front | 1.43 | 5.90 | 0.20 | 0.26 | 0.32 |
| GFC | Baseline | Interhemispheric | Front to Front | 1.21 | 5.74 | 0.27 | 0.25 | 0.31 |
| GFC | Baseline | Ipsilateral | Front to Rest | 0.06 | 5.91 | 0.96 | 0.19 | 0.19 |
| GFC | Baseline | Contralateral | Front to Rest | 0.86 | 5.25 | 0.43 | 0.18 | 0.20 |
| GFC | Baseline | Interhemispheric | Front to Rest | 1.38 | 5.57 | 0.22 | 0.18 | 0.21 |
| **Measure** | **Timepoint** | **Matrix** | **Connectivity** | **T** | **DF** | **P** | **Mean Between** | **Mean Within** |
| Frobenius | Baseline | Ipsilateral | Whole Matrix | -0.14 | 21 | 0.89 | 22.82 | 22.97 |
| Frobenius | Baseline | Contralateral | Whole Matrix | -0.25 | 19 | 0.81 | 20.66 | 20.81 |
| Frobenius | Baseline | Interhemispheric | Whole Matrix | -0.09 | 18.9 | 0.93 | 21.73 | 21.78 |
| Frobenius | Baseline | Ipsilateral | Front to Front | -0.40 | 22.33 | 0.69 | 4.70 | 4.87 |
| Frobenius | Baseline | Contralateral | Front to Front | 0.25 | 25.97 | 0.80 | 4.40 | 4.32 |
| Frobenius | Baseline | Interhemispheric | Front to Front | -0.14 | 25.67 | 0.89 | 5.00 | 5.05 |
| Frobenius | Baseline | Ipsilateral | Front to Rest | -0.20 | 22.13 | 0.84 | 9.56 | 9.67 |
| Frobenius | Baseline | Contralateral | Front to Rest | 0.11 | 25.50 | 0.92 | 8.33 | 8.30 |
| Frobenius | Baseline | Interhemispheric | Front to Rest | -0.26 | 23.69 | 0.80 | 8.92 | 9.03 |

**Supplementary Table 1. No significant baseline differences in global functional connectivity or network dissimilarity between groups.** For each measure — global functional connectivity (top panel) and Frobenius distance (bottom panel) — t-test comparisons between PFF and PBS animals at baseline are reported separately for three matrix subdivisions: the full quadrant (whole matrix: ipsilateral hemisphere, contralateral hemisphere, and interhemispheric), the frontal-to-frontal sub-matrix (within-frontal-cluster connectivity), and the frontal-to-rest sub-matrix (connectivity between the frontal cluster and all remaining regions). No comparison reached significance prior to statistical correction, indicating that the two groups did not differ systematically in functional network organization at baseline.

To quantify longitudinal changes, we computed the Frobenius distance and mean signed change (MSC) from delta connectivity matrices (early–baseline and late–baseline). LME models applied to whole-brain matrices (ipsilateral, contralateral, and interhemispheric) revealed no significant effects for the Frobenius distance (all $p$>0.05), indicating that the overall magnitude of connectivity changes did not differ between groups.

In contrast, MSC analyses revealed significant group effects for both ipsilateral ($F_{(1,6.26)} = 3.21$, $p = 0.048$) and contralateral ($F_{(1,4.75)} = 6.75$, $p = 0.051$) matrices (Supplementary Figure 10). These effects reflected a general increase in connectivity in PBS animals, particularly in the contralateral hemisphere, and a progressive decrease in PFF animals.

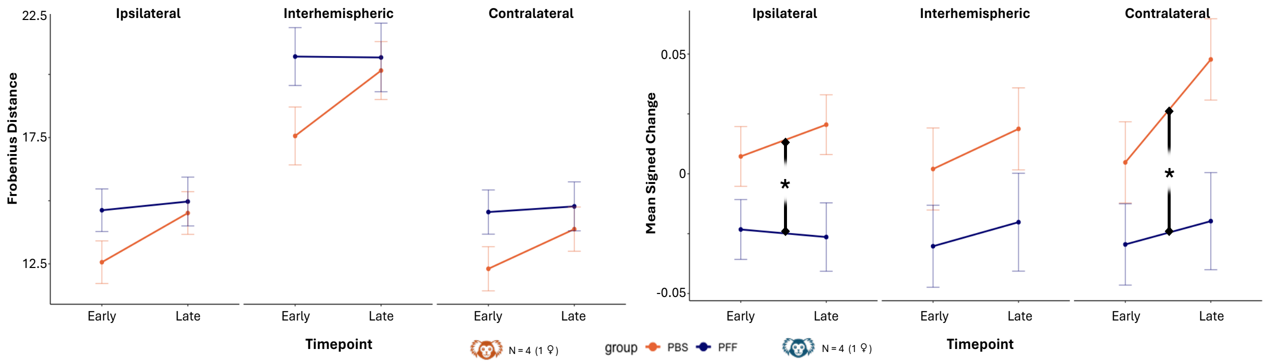


**Supplementary Figure 10. Groups differ in the direction but not the magnitude of functional connectivity change.** Frobenius distance (left panel), reflecting the overall dissimilarity between delta connectivity matrices, and Mean Signed Change (MSC, right panel), reflecting the net directional shift in functional connectivity strength, are shown for PFF (blue) and PBS (orange) animals across early (3–6 months minus baseline) and late (8–11 months minus baseline) windows, computed separately for ipsilateral, contralateral, and interhemispheric quadrants of the delta FC matrices (see Figure 4). Group means are shown as solid lines; error bars indicate the standard error of the mean. Linear mixed-effects models revealed no significant group difference in Frobenius distance, indicating comparable overall magnitudes of connectivity change between groups. A significant group effect in MSC — absent at baseline and present for ipsilateral and contralateral quadrants — was detected at both windows, indicating divergent directionality of connectivity change between PFF and PBS animals. Asterisks denote $p < 0.05$.

Given the pronounced frontal alterations observed in the connectivity matrices and their potential relevance for behavioural deficits, analyses were repeated focusing on a frontal cluster, including all regions of the frontal lobe but orbitofrontal areas in the ipsilateral and interhemispheric matrices. This cluster was selected as it exhibited the strongest decline in FC in PFF animals over time. Within this cluster, Frobenius distance did not reveal significant group differences for any matrix or comparison (all p>0.05). However, MSC analyses showed significant group effects in the ipsilateral hemisphere for both frontal-to-frontal FC ($F_{(1,11)} = 6.04$, $p = 0.032$) and frontal-to-rest connectivity ($F_{(1,6.21)} = 7.63$, $p=0.032$). For interhemispheric connectivity, a significant group effect was observed for frontal-to-rest interactions ($F_{(1,11)} = 6.37$, $p=0.029$), but not for within-frontal connectivity.

In all cases, these effects were driven by a decrease in connectivity in PFF animals relative to baseline, whereas PBS animals showed stable or mildly increased connectivity (Supplementary Figure 11).

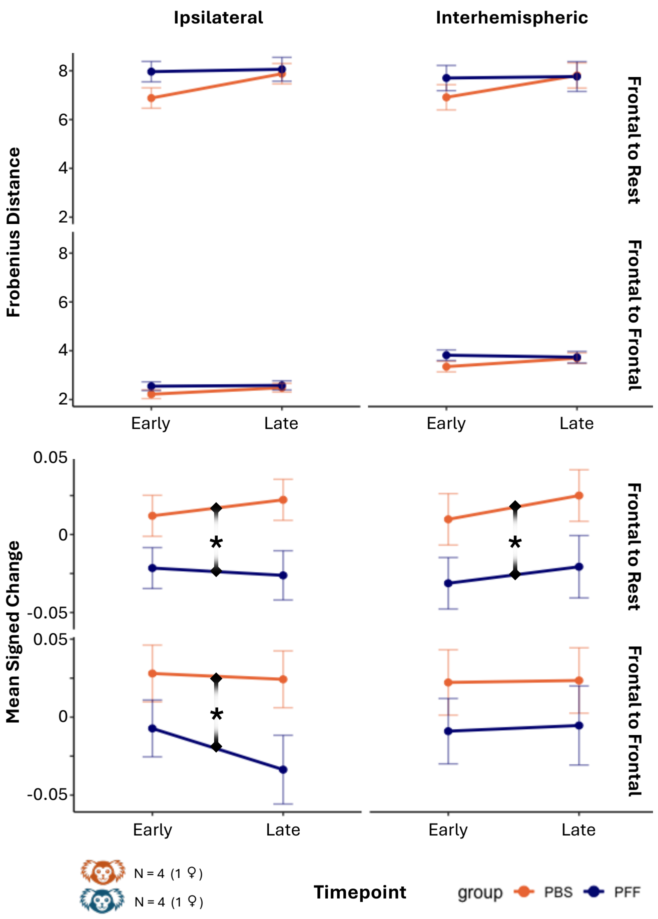


**Supplementary Figure 11. Frontal connectivity shows directional but not magnitude differences between groups, driven by frontal-to-rest connectivity.** Frobenius distance (top panel) and Mean Signed Change (MSC, bottom panel), reflecting overall matrix dissimilarity and net directional shift in functional connectivity strength respectively, are shown for PFF (blue) and PBS (orange) animals across early (3–6 months minus baseline) and late (8–11 months minus baseline) windows. Results are displayed separately for frontal-to-frontal (within-frontal-cluster connectivity, bottom row of each plot) and frontal-to-rest (connectivity between the frontal cluster and all remaining regions, top row) subdivisions, within the ipsilateral (left column) and interhemispheric (right column) quadrants of the delta FC matrices (see Figure 4 for full delta FC matrices). Group means are shown as solid lines; error bars indicate the standard error of the mean. Linear mixed-effects models revealed no significant group differences in Frobenius distance for any subdivision or quadrant. For MSC, significant group effects were detected for frontal-to-rest connectivity in both the ipsilateral and interhemispheric quadrants, and for frontal-to-frontal connectivity in the ipsilateral quadrant, with PFF animals showing a more negative delta relative to PBS animals across both windows. The group difference in interhemispheric frontal-to-frontal connectivity did not reach significance. Asterisks denote $p<0.05$.

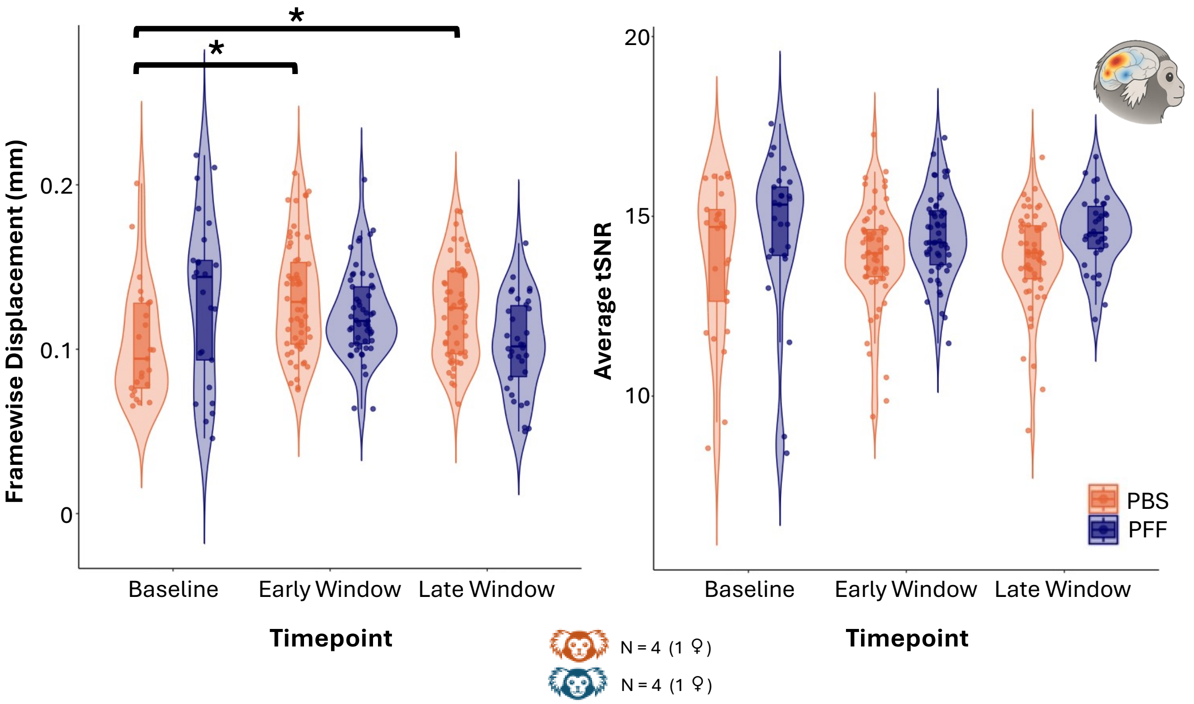


**Supplementary Figure 12. No group differences in tSNR; PBS animals show increased framewise displacement at later timepoints.** Violin plots show the distribution of average framewise displacement (FD, left panel) and temporal signal-to-noise ratio (tSNR, right panel) for PBS (orange) and PFF (blue) animals at baseline, early (3–6 months), and late (8–11 months) windows. Individual data points are overlaid. Linear mixed-effects models revealed no significant group or timepoint effects on tSNR, indicating comparable signal quality across groups and timepoints. For FD, a significant Group*Timepoint interaction was detected and followed by Holm-corrected post-hoc comparisons. The only significant difference identified was a lower FD in PBS animals at baseline relative to the PBS early and late windows; FD did not differ significantly between PBS early and late windows, nor between PFF and PBS animals at any timepoint. Asterisks denote $p < 0.05$.

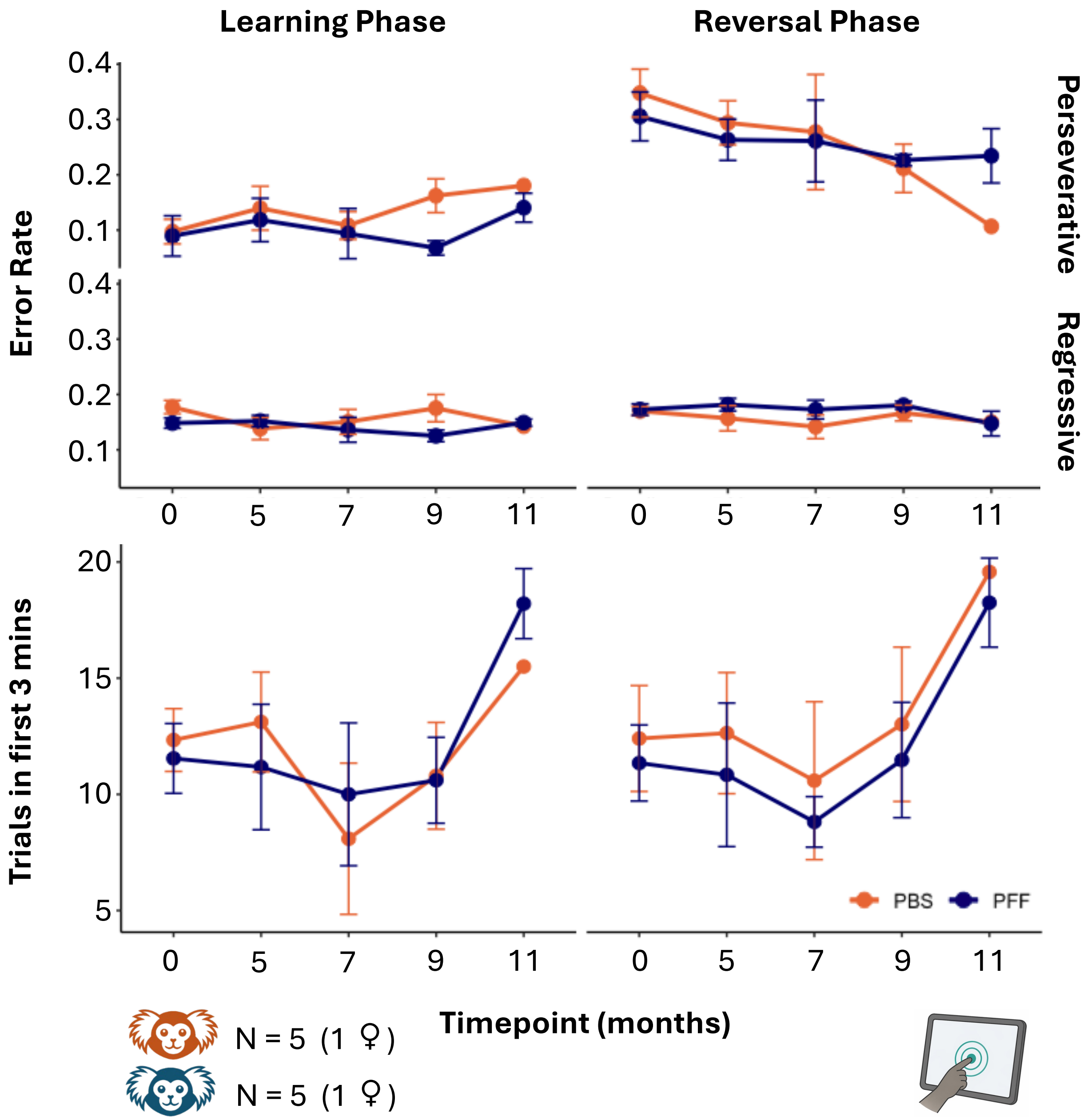


**Supplementary Figure 13. No group differences in error types or task engagement across learning and reversal phases.** Perseverative errors (first row), regressive errors (second row), and task engagement (third row; number of trials completed within the first three minutes) are shown for PBS (orange) and PFF (blue) animals across timepoints, separately for the learning phase (left column) and reversal phase (right column) of the pairwise visual discrimination task. Group means are shown as solid lines; error bars indicate the standard error of the mean. Linear mixed-effects models revealed no significant group or timepoint effects for any metric in either task phase. These results indicate that the reversal learning impairment observed in PFF animals (Figure 8) reflects a specific deficit in adaptive flexibility rather than a generalized increase in errors or reduction in task engagement.

| Marmoset ID | Site | Sex | Age at Injection (y) | Injection | fMRI | MRI | Touchscreen | Actimetry | Perfusion |
|---|---|---|---|---|---|---|---|---|---|
| Ja | Western | M | 3 | PFF | o | o | o | o | 13 months |
| Ro | Western | M | 4 | PFF | o | o | o | o | 11 months |
| Sp | Western | M | 4 | PFF | o | o | o | o | 7 months |
| Ju | Western | F | 3 | PFF | o | o |  | o | 5 months |
| Ch | Western | F | 2.5 | PFF |  |  | o |  | 7 months |
| Bu | Western | M | 2.5 | PFF |  |  | o |  | 9 months |
| Nu | Western | M | 5 | PFF |  |  |  |  | 2 months |
| Sc | Western | M | 2 | PBS | o | o | o | o | 13 months |
| Fr | Western | M | 3 | PBS | o | o | o | o | 10 months |
| Re | Western | M | 3.5 | PBS | o | o | o | o | 7 months |
| Ky | Western | F | 4.5 | PBS | o | o | o | o | 7 months |
| Ri | Western | M | 2.5 | PBS |  |  | o |  | 9 months |
| De | Western | M | 4 | PBS |  |  |  |  | 2 months |
| Am | McGill | F | 6 | PFF |  | o |  | o | 10 months |
| Da | McGill | M | 3 | PFF |  | o |  | o | 10 months |
| Si | McGill | M | 6.5 | PBS |  | o |  | o | 10 months |

**Supplementary Table 2. Subject characteristics and participation across experimental modalities.** Each row corresponds to an individual animal, identified by marmoset ID (column 1). Columns report the testing site (column 2), sex (column 3), age at injection in years (column 4), injection type — phosphate-buffered saline (PBS) or marmoset-derived αSyn preformed fibrils (PFF) — (column 5), participation in resting-state fMRI (column 6), structural MRI and deformation-based morphometry (column 7), touchscreen-based pairwise visual discrimination task (column 8), and actimetry (column 9), and time of perfusion in months post-injection (column 10). A circle in columns 6–9 indicates that the animal contributed data to the respective modality. Animals without a circle in a given column were not included in that analysis.

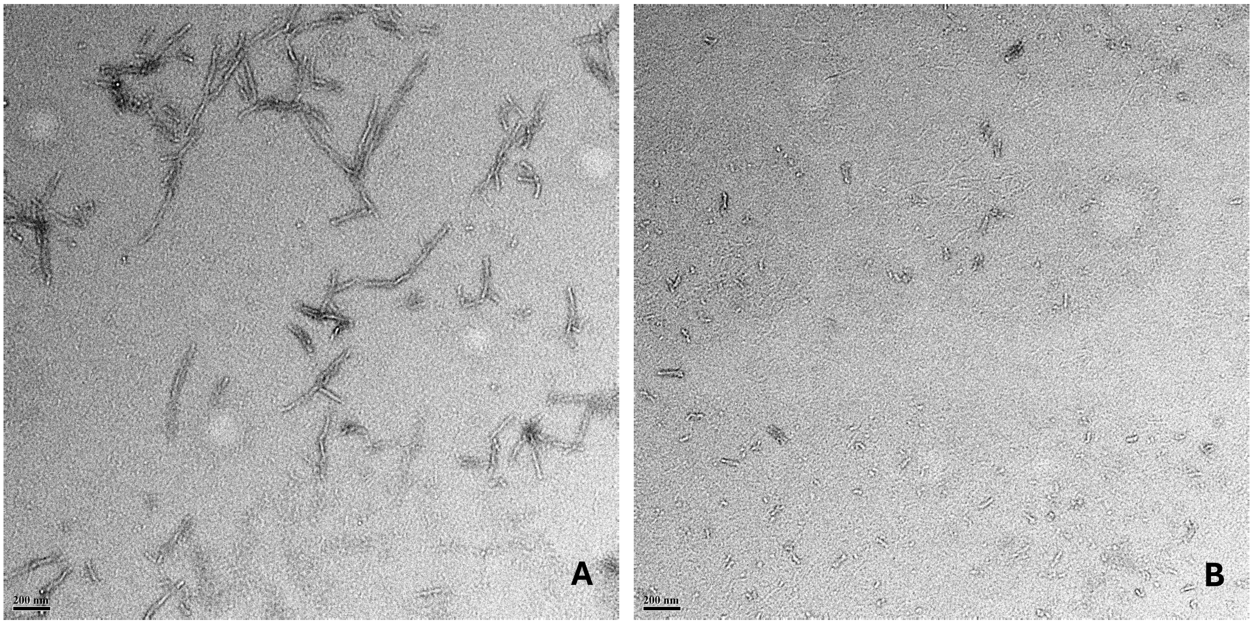


**Supplementary Figure 14. Transmission electron microscopy confirms fibril formation and fragmentation following sonication.** Negatively stained (2% uranyl acetate) transmission electron micrographs of marmoset α-synuclein preformed fibrils (PFFs) before (A) and after (B) sonication. Sonication reduced fibril length to fragments of approximately 50 nm, as confirmed by dynamic light scattering. Scale bar: 200 nm.

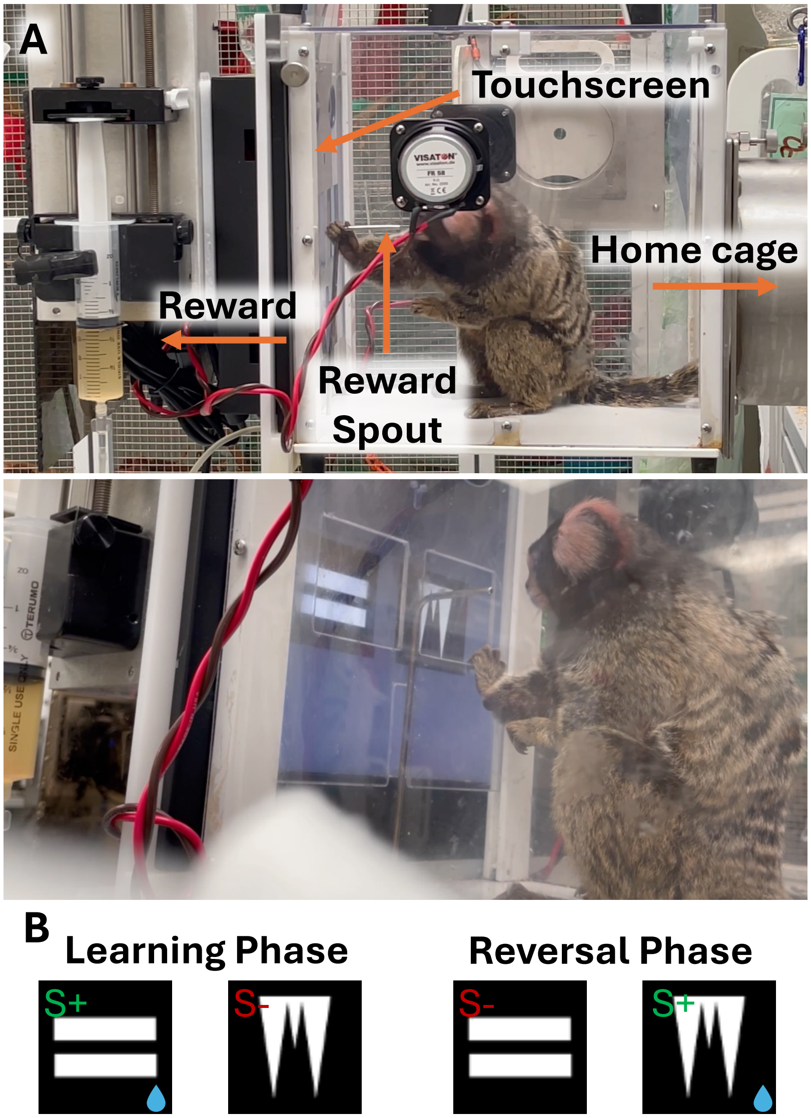


**Supplementary Figure 15. Touchscreen apparatus and task stimuli.** (A) Photographs of the touchscreen setup. The touchscreen unit is attached directly to the home cage, allowing the selected animal to be isolated within a dedicated compartment while remaining connected to the main housing area; animals were free to move between the touchscreen compartment and the isolated section of the home cage during testing. A reward spout positioned in front of the screen delivered liquid reward upon correct responses, controlled by a pump attached to the rear of the touchscreen unit. (B) Examples of visual stimuli used in the pairwise visual discrimination task. A unique pair of stimuli was used at each timepoint to prevent carry-over of stimulus-specific learning across sessions. Left: during the learning phase, one stimulus (S+) was consistently paired with reward while the other (S−) was not. Right: during the reversal phase, contingencies were reversed, such that the previously unrewarded stimulus (S−) became the rewarded one (S+) and vice versa.

206
